# Genome-wide identification of sexual-reproduction genes in fission yeast via transposon-insertion sequencing

**DOI:** 10.1101/2021.08.31.458362

**Authors:** R. Blake Billmyre, Michael T. Eickbush, Caroline J. Craig, Jeffrey J. Lange, Christopher Wood, Rachel M. Helston, Sarah E. Zanders

## Abstract

Many genes required for sexual reproduction remain to be identified. Moreover, many of the genes that are known have been characterized in distinct experiments using different conditions, which complicates understanding the relative contributions of genes to sex. To address these challenges, we developed an assay in *Schizosaccharomyces pomb*e that couples transposon mutagenesis with high-throughput sequencing (TN-seq) to quantitatively measure the fitness contribution of nonessential genes across the genome to sexual reproduction. This approach identified 532 genes that contribute to sex, including more than 200 that were not previously annotated to be involved in the process, of which more than 150 have orthologs in vertebrates. Among our verified hits was an uncharacterized gene, *ifs1* (**i**mportant **f**or **s**ex), that is required for spore viability. In two other hits, *plb1* and *alg9*, we observed a novel mutant phenotype of poor spore health wherein viable spores are produced, but the spores exhibit low fitness and are rapidly outcompeted by wildtype. Finally, we fortuitously discovered that a gene previously thought to be essential, *sdg1* (**s**ocial **d**istancing **g**ene), is instead required for growth at low cell densities. Our assay will be valuable in further studies of sexual reproduction in *S. pombe* and identifies multiple candidate genes that could contribute to sexual reproduction in other eukaryotes, including humans.

## Introduction

Sexual reproduction requires diploid cells to generate haploid gametes. Compatible gametes can then fuse to regenerate the diploid state. Thus, sexual reproduction produces recombinant offspring with the same number of chromosomes as the parents. Sexual reproduction is conserved broadly across the eukaryotic kingdom, with few exceptions [1], and thus is inferred to have been present in the common ancestor of all eukaryotes [2,3]. Long-lived asexual eukaryotic lineages are rare, suggesting that loss of sex may produce lineages that can be capable of short-term success but are unable to adapt and persist over long evolutionary time scales. As a result, asexual eukaryotic lineages have been proposed to be evolutionary dead-ends [3]. Despite the ubiquity of sexual reproduction, the set of genes required for meiosis and sexual reproduction is not fully understood even in well-studied model organisms.

Much of our understanding of the mechanisms underlying sexual reproduction was derived from classic forward genetic screens (including but not limited to [4–13]). These screens have generated enormous progress in biology but are often technically challenging because of the effort required to generate, screen, and map the mutants identified. Forward genetic screens also run the risk of incomplete saturation and thus false negatives on a whole-genome search for genes involved in a given process. Subsequently, reverse genetic screens, often of deletion collections in well-studied model organisms, have provided substantial additional insight into the meiotic toolkit of individual species (including but not limited to [14–20]). Deletion collection screens are powerful, but their use is typically limited to organisms with large research communities capable of collaboratively constructing these deletion collections. In contrast, non-model organisms rarely have these resources available, and construction of these deletion collections is cost and time prohibitive. In addition, many existing deletion collections are found in organisms that are relatively distantly related, often on the order of hundreds of millions of years diverged [21,22]. As a result, studying the evolution of rapidly evolving processes like sexual reproduction is difficult using these existing sets of tools and approaches.

An alternative high-throughput and whole genome approach is <u>T</u>ra<u>n</u>sposon insertion <u>seq</u>uencing (TN-seq)[23–26]. TN-seq works by generating massive, pooled libraries of haploid cells wherein each cell contains a single transposon insertion at a random site in the genome. These insertion sites are then specifically mapped by sequencing, often including PCR amplification of the insertion site. With sufficiently dense transposon insertions across the genome, essential genes or regions can be identified after sequence analysis as regions with significantly diminished transposon insert density relative to the rest of the genome (Figure 1A,B). Subsequent experiments can apply a selective pressure to this mutant library and then resequence to look for genes not required in the first growth condition but required in the second (Figure 1C). This approach allows pooled analysis of the equivalent of a whole genome deletion collection in a single experiment without the need to construct that entire deletion collection prior to performing the experiment. TN-seq reduces the scale of whole genome reverse genetic screens to one that can be accomplished within a single lab, rather than an entire community working together. As a result, this technique opens up the possibility of exploring gene set evolution in non-model organisms.

**Figure 1.**
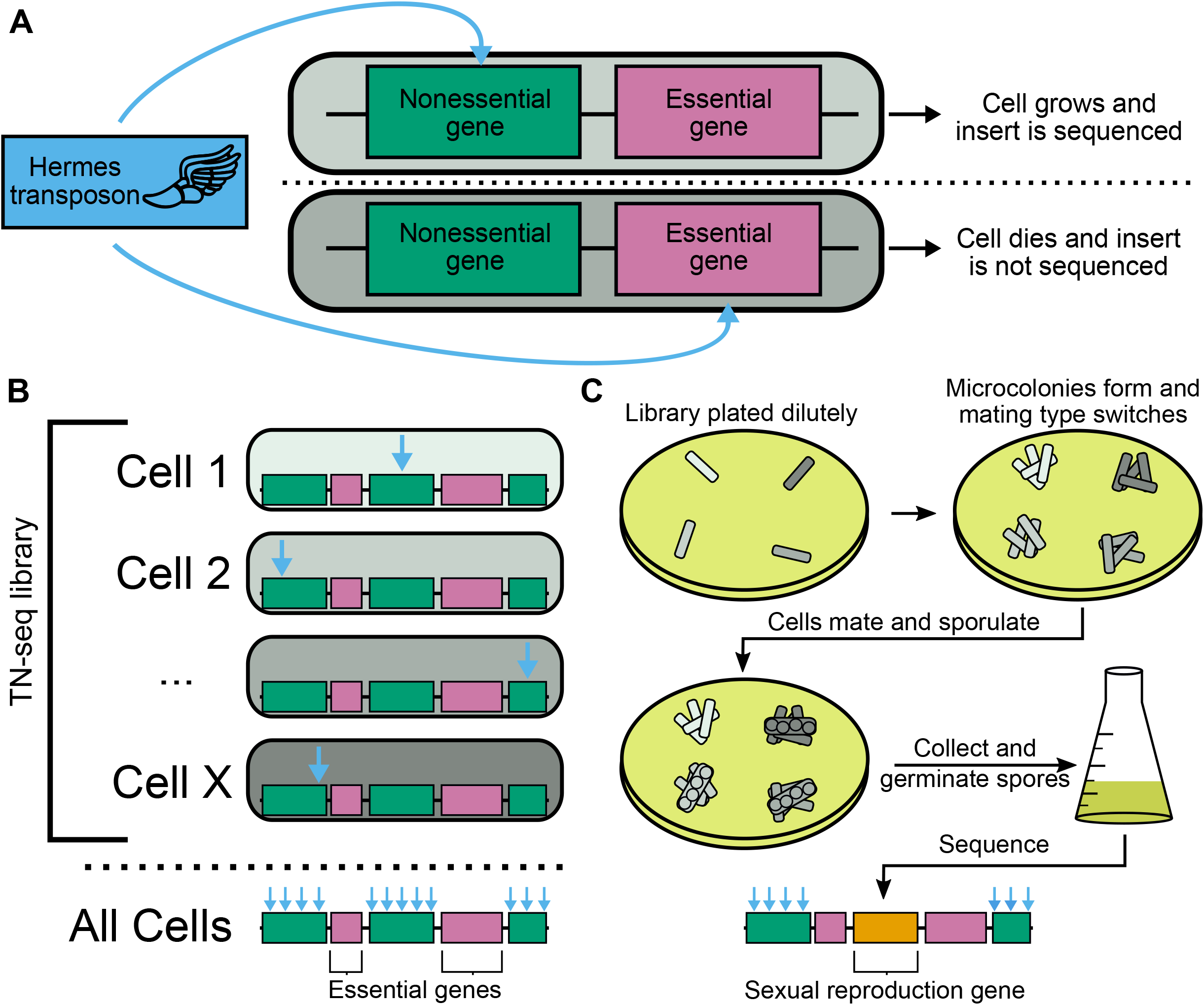
Transposon insertion sequencing can identify sexual reproduction genes. A) Transposon insertions will affect cells differently depending on where they occur in the genome. Inserts into nonessential genes (top) are unlikely to be deleterious and thus will be recovered and sequenced. In contrast, insertions into essential genes (bottom) are likely to kill the cell and not be available to sequence. B) Building a library of cells with independent transposon insertions allows mapping of essential (pink) and non-essential (green) regions of the genome by measuring insertion density via sequencing. C) A transposon insertion library can be subsequently selected for the ability to undergo sexual reproduction to produce viable spores. The library is plated at low density on MEA plates so that clonal microcolonies form. The cells within the microcolony can undergo mating type switching and mate with siblings to form diploids homozygous for a transposon insertion. The diploids then undergo meiosis and sporulation. We then collect spores, germinate them in liquid media, and assay insert frequency and location via sequencing. Genes with a role in sexual reproduction that previously tolerated insertions will no longer have inserts (orange).

TN-seq has been broadly applied in bacterial genetics where it revolutionized systems genomics [23,27]. While only used lightly in eukaryotes, it has been pioneered in fungal genetics in the model fission yeast *Schizosaccharomyces pombe* [28]. Subsequent studies in fission yeast have used TN-seq to identify factors involved in heterochromatin formation [29] and to study the fitness landscape of the genome [30]. More recently, TN-seq systems have also been developed in two other model yeasts, *Saccharomyces cerevisiae* [31,32] and *Candida albicans* [33], as well as a few less commonly studied fungi [34,35].

Here we develop and apply a screen using TN-seq to identify genes involved in sexual reproduction in the model fission yeast *S. pombe*. Previous screens have focused either on the ability to produce spores that stain appropriately with a spore dye [36] or on cytological analysis of chromosome segregation and sporulation rates [14]. This work is the first to directly measure sexual fitness consequences of mutations across the entire genome in *S. pombe*. As a result, we draw on the body of existing research on sex in *S. pombe* [14,36] to assess the validity of our TN-seq-based approach. We also identify additional genes involved in sexual reproduction that were not identified by previous approaches. For example, we identified a new class of sexual reproduction genes whose mutants exhibit competitive growth disadvantages post spore germination. Finally, we identified a mutant that failed to grow under conditions of low cell density, a condition likely to be frequently encountered by spores in nature and often employed in studies of reproduction. This work provides the basis for rapid exploration of the genes involved in sexual reproduction in numerous fungi, including other members of the *Schizosaccharomyces* genus. Further, our screen hits include many conserved genes that can be further studied in multicellular eukaryotes, in addition to *S. pombe*.

## Results

In order to determine the set of genes that contribute to sexual reproduction in *S. pombe*, we developed an assay to test the ability of cells to successfully mate, undergo meiosis, produce spores, and for those spores to germinate and grow. We took advantage of the ability of *h^90^ S. pombe* cells to switch mating types by plating a library of transposon insertion mutants at low density onto malt extract agar (MEA) plates (Figure 1C). On MEA plates, *S. pombe* cells can undergo a small number of divisions, allowing individual cells to produce microcolonies that are clonal except for the mating type locus. Once local nutrients have been exhausted, cells within these microcolonies mate to produce diploids that are homozygous for a single transposon insertion. The diploids will then quickly undergo meiosis and sporulation.

We first generated a transposon mutant library using a previously described method employing the Hermes transposon with some minor modifications (see Methods) [28]. Part of our initial mutant library was separated into aliquots for cryopreservation, while the remainder was used to prepare sequencing libraries. To begin our sexual reproduction assay, we revived and cultured one cryopreserved aliquot. Part of this culture was used to prepare a second round of pre-sex sequencing libraries, while the remainder was diluted and spread sparsely onto MEA plates. We isolated spores from the MEA plates and allowed them to germinate in liquid culture for either a “short” (single 1:100 dilution of spores into rich media and grown for 24 hours) or “long” (1:1000 dilution of spores into rich media grown for 24 hours and diluted 1:100 into rich media and grown for another 24 hours) outgrowth period. Sampling two timepoints prior to sex and two timepoints after sex allowed us to assess whether changes in insert frequencies in a given gene were specific to sexual reproduction or simply occurred continuously over time.

To identify and quantify individual transposon insertion sites at each timepoint, we amplified transposon-associated DNA via PCR and carried out Illumina sequencing with some modifications from previous protocols (see Methods)[28,30]. Importantly, our sequencing library preparation added unique barcodes to each genome fragment, as previously employed in *S. pombe* [30]. These barcodes allow us to correct for bias in PCR amplification and directly assay the frequency of a given transposon insertion in our mutant library [30].

We mapped only sequencing reads containing the PCR-amplified Hermes transposon fragments and trimmed the Hermes sequence from the reads prior to mapping. To avoid artifacts derived from repeat regions, we only used reads that mapped uniquely to the *S. pombe* genome (approximately 95% of the genome). In sum, we identified 859,876 unique transposon inserts in our original mutant library, of which we recovered 655,561 after sampling, freezing, and regrowing. After sexual reproduction, we recovered 475,815 unique inserts (72.6% of those in the plated culture) after a short outgrowth and 458,221 (69.9%) after a longer outgrowth. Insert sites present in the mutant library prior to inducing sexual reproduction can be absent from the post-sexual reproduction sets for multiple reasons. First, they may have a bona fide defect in sexual reproduction. Alternately, inserts into essential genes whose protein product has a long turnover may not be immediately eliminated from the population. In addition, mutants that failed to form microcolonies on MEA would also be expected to be depleted in our post-sexual reproduction data sets. Finally, some inserts present at low frequency in the initial set may have failed to be plated by chance and thus were not present at later time points for non-biological reasons.

In order to reduce noise from poor sampling of low frequency inserts, we filtered out all sites that originally had eight or less total inserts in our sample taken from the mutant library prior to inducing sexual reproduction. This reduced our total number of unique sites to 364,549 prior to inducing sexual reproduction, 326,269 (89.4%) in the short outgrowth post-sexual reproduction data set, and 318,729 sites in the longer outgrowth post-sexual reproduction set (87.4%). This filtering likely reduced our ability to detect true positives but also reduced the likelihood to detect false positives. However, remaining mutants still represent a high insert density across the genome with one insert every 38 bases on average.

### TN-seq identified numerous sexual reproduction candidate genes

To assess the role of a gene in sex, we measured the frequency of individual inserts in that gene within the mutant library prior to inducing sexual reproduction (pre-sex), as well as in the samples taken from germinated spores (post-sex). Inserts within many known meiotic genes displayed a clear reduction in frequency after sex, as seen for *rec12* [37,38], the ortholog of the well-studied *SPO11* endonuclease required for meiotic recombination (Figure 2A) [39]. For a more comprehensive analysis, we built a distribution of log-corrected fold changes for each gene. To do so, we set the minimum detected value for any given insert site to one read, so that sites with no detected reads in a subset of conditions were instead assumed to have one read in those conditions. By building a distribution of inserts for each gene, we were able to compare the distribution of log-corrected fold changes across a gene to that of a putatively neutral distribution using a Mann-Whitney U test. For a “neutral” control set, we analyzed inserts in intergenic regions of the genome, which had a mean log-adjusted ratio slightly below zero (i.e., no change in insert frequency between the pre- and post-sex samples) (Figure 2B). Sexual reproduction genes were defined as those which had a statistically different distribution of inserts from that of the intergenic region after Bonferroni multiple testing correction (Figure 2C).

**Figure 2.**
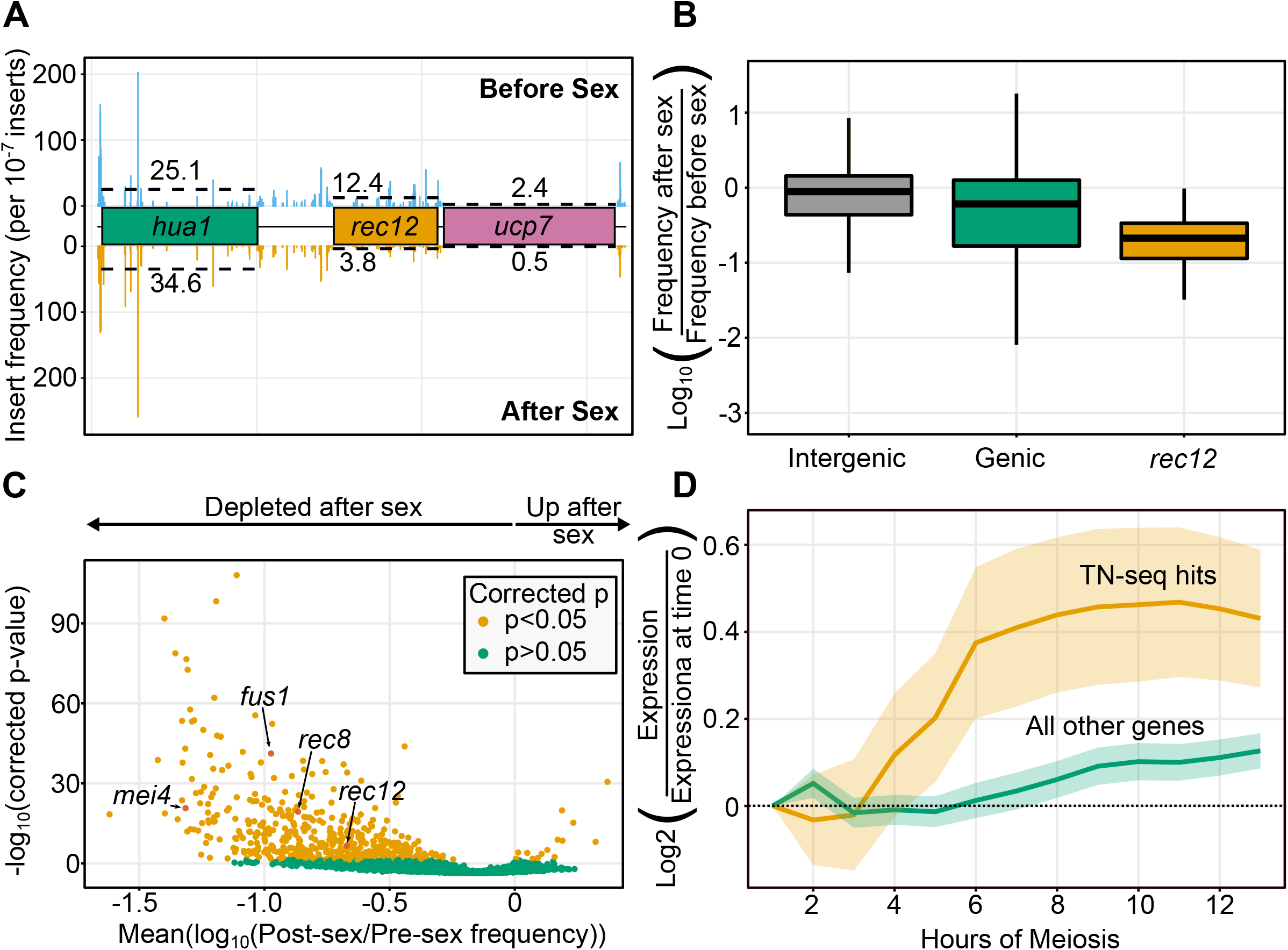
Sexual reproduction TN-seq identifies bona-fide sex genes as well as numerous candidate genes. A) Plot of transposon insert frequency across a stretch of the genome centered on the *rec12* (*SPO11*) gene. Frequencies before sex are indicated with bars above the x-axis and frequencies after sex are indicated with bars below the x-axis. Dotted line indicates mean frequency within the library of insert sites. Genes are color coded either as dispensable (green), important for sexual reproduction (orange), or essential (pink). Insert frequency is scaled to frequency per 10^−7^ inserts. B) Boxplot displaying distribution of log_10_-adjusted fold changes in insert density after sex (ie. frequency after sex/frequency before sex). Boxplots show first quartile, median, third quartile. The whiskers show the range to a maximum of 1.5 times the interquartile range above and below the first and third quartile, respectively. Outlier data points (outside the whiskers) are not displayed. This results in the removal of 5,716 of 235,578 intergenic sites, 234 of 128,971 from coding regions, and 2 of 36 from *rec12*. Inserts in intergenic regions are indicated in grey, inserts into genic regions (including introns) are shown in green, and inserts into the known meiotic gene *rec12* are shown in orange. C) Volcano plot of 3418 genes with 5 or more insert sites (out of 5118 pombe genes) displaying the mean log_10_(post-sex/pre-sex) value on the x-axis and the -log_10_ (Bonferoni corrected p-value) on the y-axis. Individual genes are shaded orange if the distribution of inserts is statistically different from the distribution of inserts into noncoding regions (p<0.05 via Mann-Whitney U test after Bonferroni correction). Genes that are not statistically different are shaded green. Four genes with known sexual reproduction defects are highlighted. D) Plot of average gene expression across meiosis for genes identified as hits with defects in sexual reproduction (orange) or not (green). Data is acquired from the Mata et al. wildtype diploid transcriptional data [42].

This approach is conservative and likely underestimates the total number of sexual reproduction genes. In addition, we did identify two categories of likely errors. First, a subset of genes had a very low number of unique insert sites and thus were difficult to score via this approach. These genes typically had either very short coding sequences or were annotated essential genes that had retained very low insert numbers at our early time points. To address this, we filtered to remove all genes which had fewer than five unique insert locations. Of the 5,118 genes annotated in *S. pombe*, 4,435 had at least one unique insert site in our library. 3,418 had at least five unique insert sites, meaning that we were able to assess function in sexual reproduction for approximately two-thirds of the pombe genome in a single pool. Of those 3,418 genes, 532 were inferred to have a role in sexual reproduction via our assay in at least one of the two outgrowth conditions, while 15 were predicted to repress sexual reproduction.

The genes predicted to repress sex revealed a second complication. Genes whose mutants have either substantial vegetative growth advantages or disadvantages will appear to have advantages or disadvantages during sex simply as a result of their vegetative growth rate. While we identified 15 genes that appeared to function as suppressors of sexual reproduction (i.e., inserts statistically increased in frequency over meiosis), most of these genes (12/15) increased in frequency at both vegetative growth steps measured as well (Figure 2 Supplement 1). Thus, most of these genes are likely not suppressors of meiosis but instead limit vegetative growth rate. As a result, we focused on genes required for meiosis and with relatively little impact on vegetative growth.

Of our 532 candidate sexual reproduction genes, 18 are *S. pombe* specific, 508 are conserved in fungi, and 373 are conserved in vertebrates. 40 of our hits lack a *S. cerevisiae* ortholog but are conserved in vertebrates. 50 hits lack an assigned name in *S. pombe*, and of those, 11 lack an ortholog in the budding yeast *S. cerevisiae* but are conserved in vertebrates [40].

Our data is consistent with previous studies of sexual reproduction in *S. pombe*. Transposon insertions into genes annotated in PomBase with decreased spore germination frequency in deletion mutants (FYPO:0000581) are significantly underrepresented in the post-meiosis dataset (p=1.2*10^−6^) [41]. Similarly, genes annotated with abnormal meiotic chromosome segregation (FYPO:0000151) are also decreased (p=2.3*10^−4^). Further, our set of genes with a function in sexual reproduction are generally more highly expressed during meiosis (Figure 2D)[42].

However, there are obvious caveats to our data as well. Numerous genes in our set of hits are required for biosynthesis of required metabolites, such as *ade2*, *ade4*, *ade6*, *ade7*, *ade8*, and *ade10*. We also have hits in genes required for arginine, asparagine, cysteine, histidine, isoleucine, leucine, lysine, methionine, proline, serine, and valine biosynthesis. These hits likely reflect a deficiency of those molecules in the MEA used for our TN-seq assay rather than a specific requirement for these genes to undergo sex and meiosis. In sum, we have 38 hits putatively involved in metabolite biosynthesis and 277 of the others have a previously annotated defect in sexual reproduction. Of the remaining 217 genes without an annotated auxotrophic or sexual reproduction phenotype, 164 are conserved in vertebrates.

### Candidate genes identified by TN-seq display sexual reproduction defects

To validate our screen we focused on four genes. Three of these genes displayed sexual reproduction defects in our TN-seq assay but lacked a relevant sexual reproduction annotation: SPAC3G6.03c (named herein *ifs1* (important for <u>s</u>ex)), *plb1*, and *alg9.* The fourth gene, *atg11*, was previously annotated with a meiotic function. *ifs1*, *plb1*, and *alg9* all had statistically significant and substantial defects in our TN-seq screen (average post-sex frequency dropped more than an order of magnitude, while *atg11* was a borderline hit that was statistically different from neutral before multiple test correction, but not after) (Figure 3A). *ifs1* encodes a maf-like protein that has not been explored in *S. pombe*, but homologs in bacteria act to inhibit septation [43]. *plb1* encodes a phospholipase B enzyme, which is involved in lipid metabolism. Mutants of *plb1* exhibit sensitivity to osmotic stress as well as loss of nutrient repression of mating at 20°C, resulting in an increase in mating efficiency rather than the poor fitness observed in our assay [44]. *alg9* encodes an alpha-1,2-mannosyltransferase that has no obvious link to meiosis aside from one report of a weak physical association with *moc3* and *moc1*/*sds23*, which are important for sexual differentiation [45].

**Figure 3.**
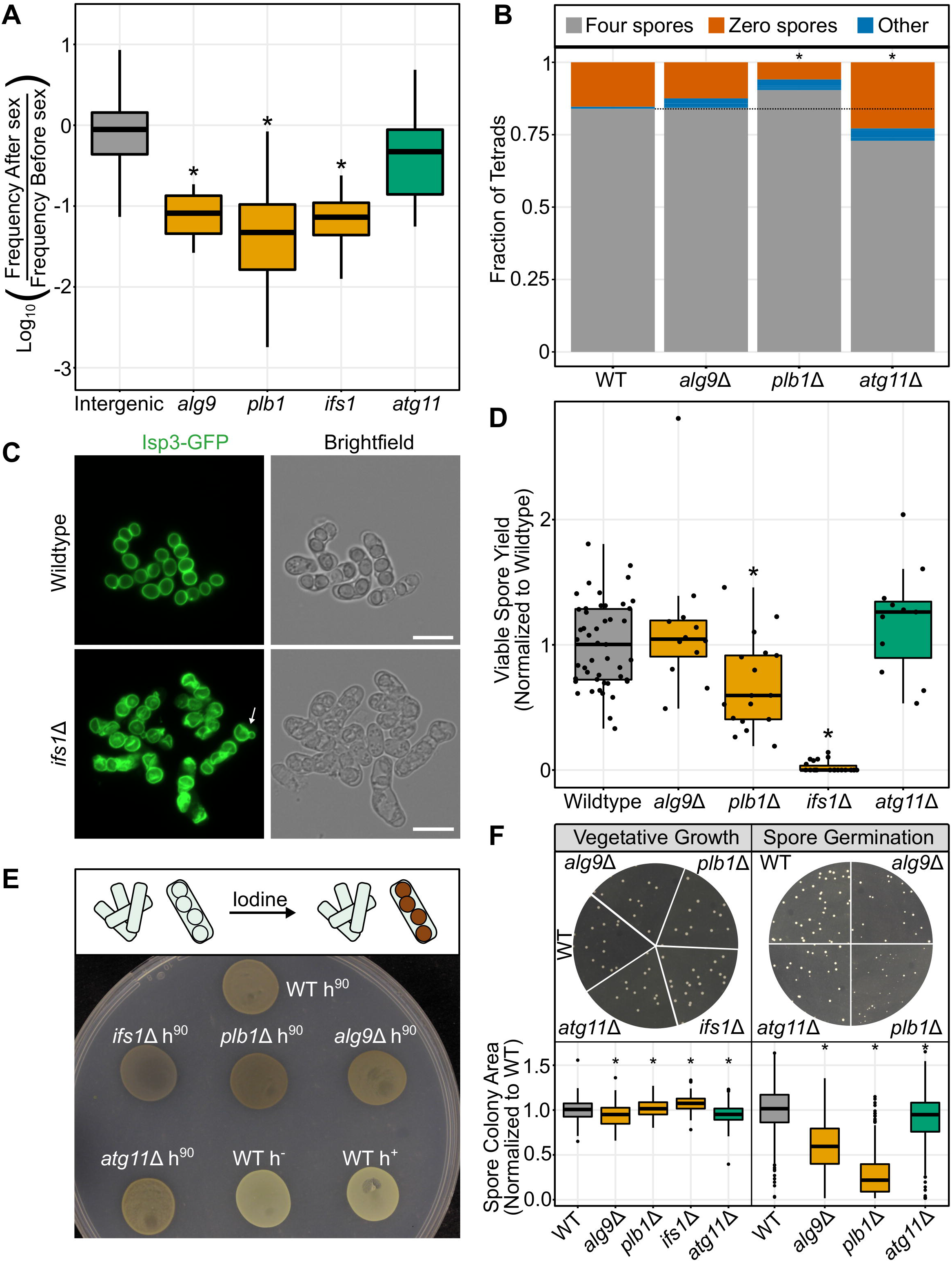
TN-seq identified candidates have defects in spore germination and spore health. A) Boxplot displaying distribution of log_10_-adjusted fold changes in insert density after sex (ie. frequency after sex/frequency before sex). Boxplots show first quartile, median, third quartile. The whiskers show the range to a maximum of 1.5 times the interquartile range above and below the first and third quartile, respectively. Outlier data points (outside the whiskers) are not displayed. This results in the removal of 5,716 of 235,578 intergenic sites, 2 of 16 sites from *alg9*, 0 of 166 sites from *plb1*, 2 of 38 sites from *ifs1* and 0 of 25 from *atg11*. Inserts in intergenic regions are indicated in grey, genes where inserts were significantly depleted after sexual reproduction in the TN-seq data are shown in orange (*alg9*, *plb1*, *ifs1*) and genes where inserts were not significantly depleted after sexual reproduction are shown in green (*atg11*). B) Scoring of acsi produced by the indicated mutants and wildtype. Asci were scored as having either four spores, no visible spores, or any other category (1, 2, 3, or more than 4). *ifs1*Δ mutants were not scored because the spores were atypical in appearance and difficult to accurately identify. Fisher’s exact test, WT, n=287; *alg9*Δ, n=337, p=1.0; *plb1*Δ, n=271, p=0.031; *atg11*Δ, n=491, p=0.0004. C) Isp3-GFP was visualized in wildtype and *ifs1*Δ mutants on an AXIO Observer.Z1 (Zeiss) wide-field microscope with a 40x C-Apochromat (1.2 NA) water-immersion objective after incubation for 2 days at 25°C on an MEA plate. Scale bars indicate 10 microns. Arrow indicates “snowman” spore. D) Viable spore yield assay showing on the y-axis the number of spores produced per yeast cell plated, normalized to the mean value for wildtype. All mutants were assayed in a set of at least 5 biological replicates alongside at least 5 wildtype replicates. Points display results from a single replicate, normalized to the mean from the corresponding wildtype controls. The boxplots summarize the underlying points and show first quartile, median, third quartile while the whiskers show the range of the data to a maximum of 1.5 times the interquartile range below and above the first and third quartile, respectively. Points outside the whiskers can be considered outliers. Mann-Whitney U Test, *alg9*Δ, p =0.61; *plb1*Δ, p =0.0028; *ifs1*Δ, p= 1.18* 10^−11^; *atg11*Δ, p = 0.21. E) Iodine staining of *h^90^* mutants after 3 days on SPAS media at 25°C temperature. Iodine should stain spores brown, while leaving unsporulated cells unstained (top). Wildtype *h^90^* is shown as an iodine-staining positive control and wildtype *h*^−^ and *h*^+^ strains are shown as non-staining negative controls. F) Plates illustrating colony size for wildtype and *plb1*Δ mutant strains after 3 days of growth at 32°C on YEA+sup plates. Vegetative growth was plated from a serial dilution of an overnight culture in YEL+sup at 32°C. Spore growth was plated from a serial dilution of glusulase-treated spores that were originally generated after 3 days on MEA at 25°C. All spores were generated, plated, and imaged at the same time. The bottom plot displays quantification of colony sizes using Fiji to perform thresholding and particle analysis. The boxplots summarize measurements from 4-5 separate plate images for each genotype and show first quartile, median, third quartile while the whiskers show the range of the data to a maximum of 1.5 times the interquartile range below and above the first and third quartile, respectively. Points outside the whiskers can be considered outliers. All four mutants produced yeast colonies that were on average statistically significantly different in size from wildtype (Mann-Whitney U test, wildtype, n=181; *alg9*Δ, n= 183, p=4.3 * 10^−6^; *plb1*Δ, n=213, p= 0.033; *ifs1*Δ, n=210, p = 2.8 * 10^−12^; *atg11*Δ, n=164 p=0.00034) All three mutants tested produced spore colonies that were statistically significantly smaller than wildtype (Mann-Whitney U test, wildtype, n=467; *alg9*Δ, n=213, p<2.2 * 10^−16^; *plb1*Δ, n=388, p= <2.2 * 10^−16^; *atg11*Δ, n=286, p=3.3 * 10^−6^).

In order to test the hypothesis that these genes are important for sexual reproduction, we began by individually deleting each gene in a common strain background. Mutants of all four genes successfully mated and formed asci on MEA, although *plb1*Δ mutants produced asci that were unusually flocculant (i.e., clumpy) (Figure 3 Supplement 1). We quantified the frequency of typical four-spore tetrads for each mutant (Figure 3B). Both *plb1*Δ and *atg11*Δ were significantly different from wildtype (Fisher’s exact test, WT, n=287; *alg9*Δ, n=337, p=1.0; *plb1*Δ, n=271, p=0.031; *atg11*Δ, n=491, p=0.0004), although in opposite directions, with *plb1*Δ mutants producing a slightly higher percentage of four-spore tetrads and *atg11*Δ mutants producing slightly less.

*ifs1*Δ mutant asci were difficult to score via brightfield imaging because spores did not appear to properly individualize, although vaguely spore-like shapes were sometimes visible within asci (Figure 3 Supplement 1). To better visualize *ifs1*Δ mutant spores, we generated a deletion of *ifs1* in an *isp3*-GFP strain. Isp3 coats the exterior of wildtype spores (Figure 3C, top) [46]. Imaging Isp3-GFP in asci generated on MEA media revealed that *ifs1*Δ mutants made wrinkled, irregular spores that frequently had blebs on the exterior, reminiscent of previously described “snowman” spores (Figure 3C) [47]. This phenotype was less severe for asci generated on SPA media, where spores were still frequently snowman-shaped, but otherwise had less severe structural defects (Figure 3 Supplement 2). Because imaging asci generated on MEA required scraping cells off of plates to image, while spores could be imaged in place on SPA, it is possible that *ifs1*Δ spores are more fragile than wildtype spores and the more severe phenotype on MEA is simply the result of increased manipulation for imaging.

We next used a viable spore yield (VSY) assay to assess fertility. Briefly, viable spore yield measures the total number of viable spores produced per cell plated on the original mating plates [48]. Two mutants (*ifs1*Δ and *plb1*Δ) had statistically decreased viable spore production via this assay after three days incubating on MEA mating plates in dense cell patches (Mann-Whitney U Test, *alg9*Δ, p =0.61; *plb1*Δ, p =0.0028; *ifs1*Δ, p= 1.18* 10^−11^; *atg11*Δ, p = 0.21) (Figure 3D). The phenotype of *ifs1*Δ was very strong. Spores very rarely germinated to form colonies, which was unsurprising given the extreme spore formation defects we observed. We retested two *ifs1*Δ spore colonies via viable spore yield to test the hypothesis that a rare genetic suppressor had arisen in these spores that rescued the *ifs1*Δ mutant phenotype. However, these survivors retained the mutant phenotype, suggesting instead that the function of *ifs1*, while extremely important, is not absolutely required for spore germination (Figure 3 Supplement 3).

Previous studies of *S. pombe* have utilized the fact that sexual spores will stain brown when exposed to iodine while vegetative cells will not in order to rapidly screen for mutants with defects in sexual reproduction (Figure 3E, top) [13,36,49,50]. This phenotype is visible without magnification in colonies on mating plates so it is simple to screen large numbers of colonies. Because we knew *ifs1*Δ mutants produced very few viable spores, we tested whether all of our mutants produced patches that stained with iodine. All four of our tested mutants stained with iodine on mating media, indicating that even though *ifs1*Δ spores almost never germinate and appear atypical via microscopy, they still stain with iodine (Figure 3E, bottom). This may help explain why some of our TN-seq hits were missed in previous screens using iodine to look for sporulation defects in *S. pombe*.

### *plb1* mutants exhibit a gamete health defect

While the defect in the *ifs1*Δ mutant was nearly complete, *plb1*Δ mutants had a much more subtle defect via the viable spore yield assay and both *atg11*Δ and *alg9*Δ mutants appeared to have no defect at all. Because our TN-seq data led us to expect a defect in viable spore yield for *plb1*Δ and *alg9*Δ, we assayed these mutants under other conditions, but did not reproduce the strong defect in reproductive fitness for either mutant detected in our TN-Seq assay (Figure 3 Supplement 4, A-D). In fact, reproducing the conditions originally used in the TN-seq assay for the viable spore yield assay generated a small *increase* in viable spore yield for *alg9*Δ (Mann-Whitney U Test, *alg9*Δ, p=0.0022; *plb1*Δ, p= 0.24; Figure 3 Supplement 4D). Combining all the data generated for these mutants in this study across multiple conditions resulted in no distinguishable defect in viable spore yield relative to wildtype (Mann-Whitney U Test, *alg9*Δ, p=0.0013 with higher mean than wildtype; *plb1*Δ, p= 0.73) (Figure 3 Supplement 4E).

However, we noticed that colonies produced by germinating *plb1*Δ mutant spores were variable in size and were often substantially smaller than those produced by wildtype spores (Figure 3F, top right). Colonies derived from *alg9*Δ mutant spores similarly produced visually smaller colonies than wildtype spores. To assess this quantitatively, we imaged colonies generated from spores for wildtype, *alg9*Δ, *plb1*Δ, and *atg11*Δ mutants and measured colony size computationally. As expected, *alg9*Δ and *plb1*Δ mutants produced smaller colonies, but surprisingly, *atg11*Δ mutant spores also had a relatively subtle growth defect as well (Mann-Whitney U test, p=3.3 * 10^−6^) (Figure 3F, bottom). As a control, we performed the same assay by plating vegetatively growing cells and measuring colony sizes. All four mutants tested were significantly different from wildtype, but with substantially smaller magnitudes of change. Notably, *ifs1*Δ and *plb1*Δ mutants made slightly larger colonies than the wildtype on average.

As a result, we predicted that when grown in competition with wildtype spores, *plb1*Δ, *alg9*Δ, and *atg11*Δ mutant spores would have a far more dramatic fitness disadvantage during spore germination resulting from either delayed germination or slow growth following spore germination. In order to test this hypothesis, we performed a competitive growth assay (Figure 4A). We mixed approximately equal ratios of wildtype and mutant spores in rich liquid media to allow them to germinate and grow. We took an initial sample to verify that we had properly mixed our spores. Then we allowed growth for 24 hours and plated again. We assessed the frequency of the two genotypes by replica plating colonies to media containing G418. Wildtype cells die on this media, but each of our mutants survive because we replaced the target coding sequence with the kanMX4 gene. As a control, we also performed the same assay by competing a presumably neutral deletion of a *wtf12* pseudogene marked with the same selective marker [51] against the same wildtype strain. This assay revealed that both *plb1*Δ and *alg9*Δ mutant spores had a substantial competitive growth defect, while *atg11*Δ mutant spores once again had a relatively subtle defect (Mann-Whitney U test, *alg9*Δ, p= 1.9*10^−10^; *plb1*Δ, p=1.5*10^−8^; *atg11*Δ, p=0.0483) (Figure 4B, right). We also performed the same assay starting with vegetatively growing yeast instead of sexual spores. Unlike spore germination, *plb1*Δ and *atg11*Δ mutants were not defective in competitive vegetative growth in our assay, suggesting a defect specifically during spore germination (Figure 4B, left). In contrast, *alg9*Δ mutants did exhibit a vegetative growth defect in this assay, although the magnitude was much smaller than that observed for spores and was not sufficient to explain the defect in spore growth (Mann-Whitney U test, *alg9*Δ p= 3.3*10^−7^; *plb1*Δ, p=0.17; *atg11*Δ, p=0.8723).

**Figure 4.**
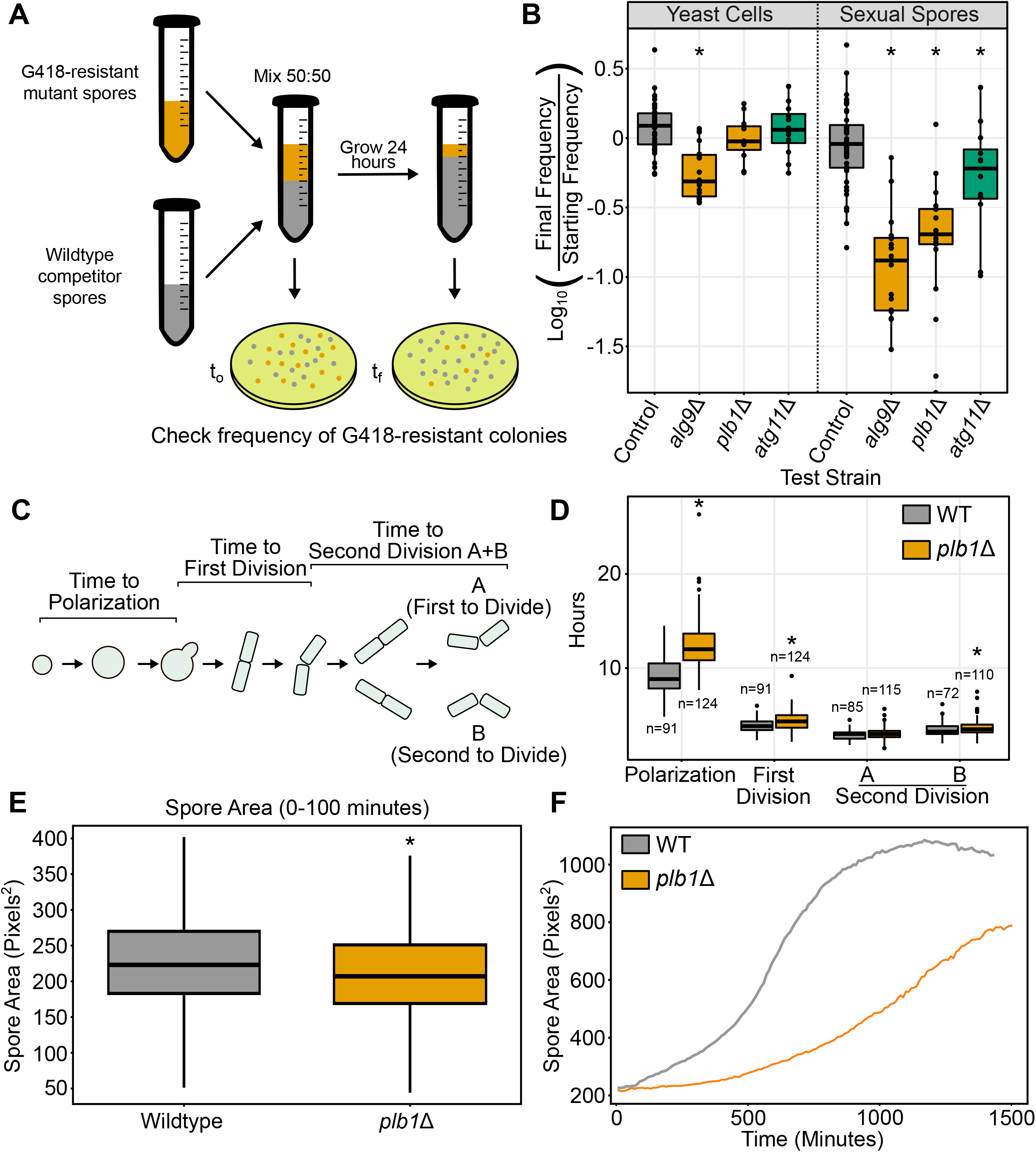
*plb1*Δ mutant spores germinate more slowly in a size-dependent fashion. A) Schematic showing spore competition assay. Spores are mixed at a 50:50 ratio and incubated with shaking overnight at 32°C in rich media (YEL). Samples of starting and final cultures are plated to YEA+sup media. Once colonies grow, they are replicated to YEA+sup with G418 and the proportion of G418-resistant colonies is counted. B) Results of spore and yeast competition experiments. Results are a log ratio of the final frequency normalized to the initial frequency. Points indicate results of individual replicates from two experiments, each including at least 5 biological replicates. The boxplots summarize the underlying points and shows first quartile, median, third quartile while the whiskers show the range of the data to a maximum of 1.5 times the interquartile range below and above the first and third quartile, respectively. Points outside the whiskers can be considered outliers. C) *S. pombe* spores undergo several landmarks in the process of germinating. Spores initially begin growing isotropically (ie. swelling). This phase ends when cells begin polarized growth and elongate on one side. This cell will eventually divide by fission for the first time and each of those daughters will go on to divide a second time after some delay. We scored each of these landmarks manually from videos using Fiji. D) Videos of spore germination were scored and time between each step in spore germination was tracked for individual spores. The boxplot shows first quartile, median, and third quartile while the whiskers show the range of the data to a maximum of 1.5 times the interquartile range below and above the first and third quartile, respectively. Points outside the whiskers can be considered outliers. E) Boxplot of spore sizes from the first 10 frames (100 minutes) of videos of spore germination. Mutant and wildtype are each derived from at least two videos each from two separate days. Spores were identified using deep learning (see methods). Boxplot displays first quartile, median, and second quartile, while the whiskers show the range of the data to a maximum of 1.5 times the interquartile range below and above the first and third quartile, respectively (N= 53,841 spores for wildtype and 73,448 spores for *plb1*Δ mutants; p= 2.45 * 10^−88^, T-test) F) Plot of average spore area over the course of videos of spore germination. Spores were identified via deep learning. Data are derived from at least two videos each from two separate days. Average spore sizes stabilize once spores begin to divide and grow vegetatively as yeast cells. Data are truncated at 1500 minutes as dividing cells begin to affect the average in *plb1*Δ mutants. The average size of *plb1*Δ mutant cells does not reach wildtype levels by the time cells have divided enough to make continued tracking impossible with this approach.

Defects in chromosome segregation often result in both a reduction in frequency of viable spores and production of viable but slow growing aneuploid gametes. Because *plb1*Δ and *alg9*Δ mutants showed a moderate decrease in spore viability and a decrease in competitive fitness, we explored the hypothesis that these mutants produced aneuploids at a higher rate than wildtype. To do so, we introduced a codominant marker system for chromosome III. In this system, one parent strain contains a wildtype *ade6* locus while the other strain of the opposite mating type contains an *ade6* locus that has been deleted with the *hphMX6* marker [52]. As a result, progeny can only be both Ade+ and hygromycin resistant if they carry both copies of this allele, suggesting chromosome III disomy. Chromosome III is the only viable single gain of chromosome aneuploidy in *S. pombe* [53]. This assay revealed that neither *plb1*Δ nor *alg9*Δ mutants exhibit elevated levels of disomy for chromosome III (Table 1), suggesting that their slow growth defects are instead the result of a distinct defect in spore fitness.

**Table 1.**

|  | #euploid | #disomes III | % disomic III |  |
| --- | --- | --- | --- | --- |
| *plb1Δ* | 535 | 19 | 3.4% | p=0.4829<br>Fisher's Exact<br>Test |
| Wildtype | 796 | 35 | 4.2% |  |
| *alg9Δ* | 144 | 2 | 1.4% | p=0.1103<br>Fisher's Exact<br>Test |
| Wildtype | 157 | 8 | 4.8% |  |

To further examine the spore fitness defect we observed, we decided to focus on *plb1*Δ mutants and performed time lapse imaging of spore germination. Briefly, we plated either wildtype or *plb1*Δ mutant spores densely onto an agar plate. Immediately after the plates dried, we took a punch from the plate, placed it onto a coverslip, and began continuous imaging on a wide field scope at 32°C for 24-48 hours. We began by manually scoring several landmarks during *S. pombe* spore germination (Figure 4C). This revealed a dramatic difference in initial growth rate following spore germination (Figure 4D). *plb1*Δ mutant spores took much longer than wildtype spores to first establish polarity as well as to undergo the first mitotic division once they had established polarity than wildtype spores. However, most of the delay was prior to polarization, and by the second division, only the later of the two daughter divisions was still significantly slower (Mann-Whitney U test, polarization, p< 2.2 * 10^−16^; first division, p=0.00072; second division A, p= 0.2029; second division B, p= 0.026).

### *plb1*Δ mutant spores are smaller and germinate more slowly than wildtype spores

In order to extract growth parameters from these videos, we implemented a deep learning approach to identify spores and track them throughout our imaging. This approach revealed that *plb1*Δ mutant spores were on average smaller than those observed in wildtype (T-test, p= 2.45 * 10^−88^; wildtype, n=53,841; *plb1*Δ, n=73,448) (Figure 4E). Further, monitoring spore size throughout the videos via deep learning revealed that *plb1*Δ mutant spores on average grew more slowly than wildtype spores (Figure 4F). Extracting growth parameters from all identified spores recapitulated what we had seen via our manual tracking, where spores first grew symmetrically in all directions before eventually establishing polarity (Figure 4 Supplement 1, Supplemental Video 1). The length of our videos was limited by crowding of dividing cells and ended well before visible colonies would form to be counted in a viable spore yield assay (1-2 days versus 5 days however, a significant portion of *plb1*Δ mutant spores had not visibly grown by the end of the videos, despite yielding a near normal viable spore yield. Our data demonstrate that spores produced by *plb1*Δ are generally viable but exhibit substantial delays (i.e., more than 24 hours) in germination and this is the origin of the fitness defect in sexual reproduction detected in our TN-seq assay. Taken together, our data suggest that in wildtype cells Plb1 contributes to efficient spore germination and progression to cell division.

### *sdg1* is required for low density growth

Interestingly, our TN-seq analysis also identified 4 genes (*sfb3*, *ggt1*, SPAC12B10.02c, SPAC18B11.11) as significant contributors to sexual reproduction that are currently annotated as essential (Figure 5A) [54]. This was curious because our assay should only be able to query non-essential genes in which transposon insertions are present in our pre-sex sample. We attempted to delete both *ggt1* and *SPAC12B10.02c* but only successfully obtained mutants of *SPAC12B10.02c*. The *SPAC12B10.02c*Δ mutants demonstrated an unexpected “unstreakable” phenotype where cells grew at the start of a streak where the initial cell density was high, but they failed to robustly grow to a visible colony from the portions of a streak where single colonies should be present (Figure 5B). We confirmed this phenotype more formally via a spot dilution assay, where cells grew similarly to wildtype at higher cell densities and failed to grow as the initial concentration of cells plated became more dilute (Figure 5C). Because this gene appears to be required for individual cells to survive and grow without other cells nearby, we named this gene *sdg1* for <u>S</u>ocial <u>D</u>istancing <u>G</u>ene. The ability to grow at low density is likely an important trait for yeast both in nature and is also important to understand in the context of laboratory analysis of sexual reproduction. Many basic assays for studying sex, such as tetrad dissection or even plating colony forming units for a viable spore assay, require cells to be plated at low densities and assume that a strain will behave similarly, independent of plating density. *sdg1*Δ mutants clearly violate these assumptions.

**Figure 5.**
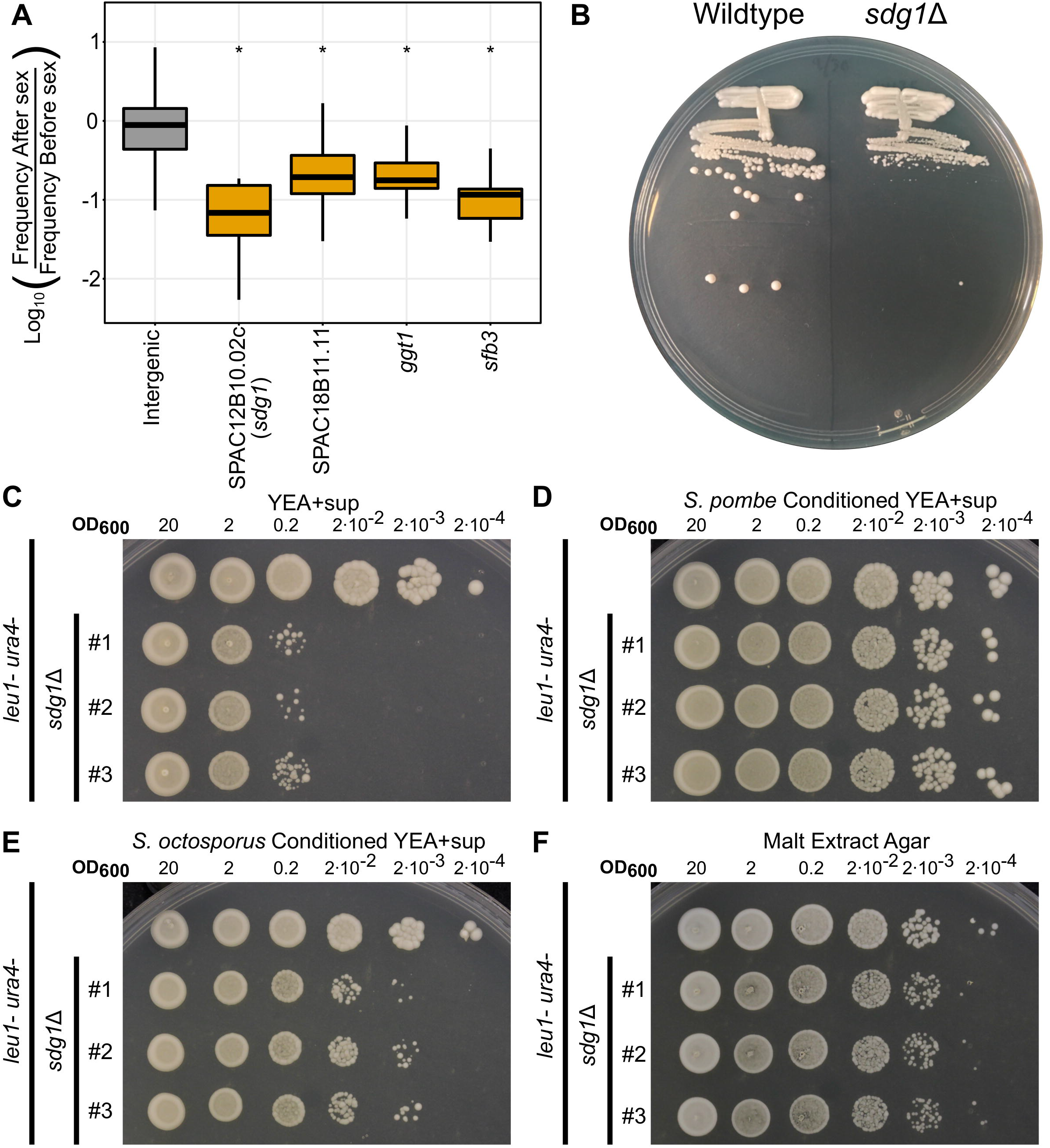
*sdg1*Δ mutants have a density-dependent growth defect that can be rescued by conditioned media. A) Boxplot displaying distribution of log_10_-adjusted fold changes in insert density after sex (ie. frequency after sex/frequency before sex). Boxplots show first quartile, median, third quartile. The whiskers show the range to a maximum of 1.5 times the interquartile range above and below the first and third quartile, respectively. Outlier data points (outside the whiskers) are not displayed. This results in the removal of 5,716 of 235,578 intergenic sites, 0 of 24 from SPAC12B10.02c/*sdg1*, 3 of 148 from SPAC18B11.11, 2 of 22 from *ggt1*, and 1 of 14 from *sfb3*. Inserts in noncoding regions are indicated in grey and inserts into candidate genes are shown in orange. B) Streaking assay showing a parent *S. pombe* strain (*ura4*-D18, *leu1*-32) and an *sdg1*Δ mutant (*sdg1*Δ::kanMX4, *ura4*-D18, *leu1*-32) struck to single colonies on a YEA+sup plate. C-F) Spot dilution assays with 5 μL spots plated. The initial leftmost spot is of OD600 = 20 culture and each successive spot is a 10-fold dilution, so that the final spot should be 10^5^ less concentrated than the first. All four experiments were conducted on the same day with the same dilution series of parent strain (*ura4*-D18, *leu1*-32) and three independent *sdg1*Δ mutants on the same genetic background (*sdg1*Δ::kanMX4, *ura4*-D18, *leu1*-32). C) Spotted to standard yeast extract agar with supplements (YEA+sup) and incubated for 4 days at 32°C. D) Spotted to conditioned yeast extract agar + supplements (see methods) made from parent strain *S. pombe* (*ura4*-D18, *leu1*-32) and incubated for 4 days at 32°C. E) Spotted to conditioned yeast extract agar made from wildtype *S. octosporus* and incubated for 4 days at 32°C. F) Spotted to malt extract agar and incubated at 25°C for 4 days.

Very little is currently known about *sdg1* in *S. pombe*. PomBase describes it as an ortholog of *PHO86* in *S. cerevisiae*, which encodes an ER resident protein required for protein packaging into COPII vesicles [54]. However, *PHO86* and *sdg1* are only very distantly related, if at all. Three progressive rounds of searches using JACKHMMER [55], a hidden Markov Model based protein homology tool, starting from either *sdg1* in *S. pombe* or *PHO86* in *S. cerevisiae* fail to identify the other protein. JACKHMMER searches starting from *PHO86* do eventually identify other fungal proteins with the same N-acyltransferase domain as *sdg1*, including a protein from *Schizosaccharomyces cryophilus* [56] that is an ortholog of *S. pombe naa50*, rather than *sdg1*.

The density-dependent growth phenotype of *sdg1*Δ mutants was consistent with a quorum sensing effect where cell growth is dependent on factors secreted by neighboring cells. To test this hypothesis, we generated conditioned YEA+sup (yeast extract agar with supplements) media by replacing half of the water typically in the plates with filtered conditioned YEL+sup (yeast extract liquid with supplements) media previously used to grow *S. pombe* cells to saturation. We found that conditioned media largely rescues the low-density growth defect of *sdg1*Δ mutants (Figure 5D), while simply adding more unconditioned media did not (Figure 5 Supplement 1A, B). Media conditioned with *sdg1*Δ mutant cells conferred a similar rescue, suggesting that Sdg1 is not itself important for production of a quorum sensing signal (Figure 5 Supplement 1C, D). These experiments also suggest that the signal responsible for rescuing low density growth survives autoclaving.

In addition, generating plates using media conditioned by *Schizosaccharomyces octosporus* growth confers intermediate rescue of the low-density growth defect (Figure 5E, Figure 5 Supplement 1E), while media conditioned by the more distant relative *Schizosaccharomyces japonicus* provided minimal rescue (Figure 5 Supplement 1F). Surprisingly, media conditioned by the budding yeast *Saccharomyces cerevisiae* provided nearly the same level of rescue as *S. octosporus* (Figure 5 Supplement 1G). This suggests that the quorum sensing signal is only partially conserved between these species.

Surprisingly, the low-density growth defect of *sdg1*Δ mutants was largely suppressed on MEA plates (Figure 5F). While *sdg1*Δ colonies still grew more slowly on MEA plates than on standard YEA+sup plates, they grew at similar plated cell densities to wildtype cells, which was not true when plated on YEA+sup. We hypothesized that *sdg1*Δ mutants had performed poorly in our TN-seq assay because we plated at low density to acquire clonal microcolonies rather than because they had a bona fide sexual reproduction defect.

However, testing viable spore yield was complicated by the fact that *sdg1*Δ mutant spores fail to grow when plated at low enough densities to count colonies accurately (Figure 6A). As a result, we employed a modified version of our assay where we plated spores to conditioned YEA+sup media instead of standard YEA+sup. This did not affect the viable spore yield of wildtype *S. pombe* (Figure 6B, left). This assay revealed that *sdg1*Δ mutants did not significantly affect viable spore yield (Mann-Whitney U test, p=0.9225).

**Figure 6.**
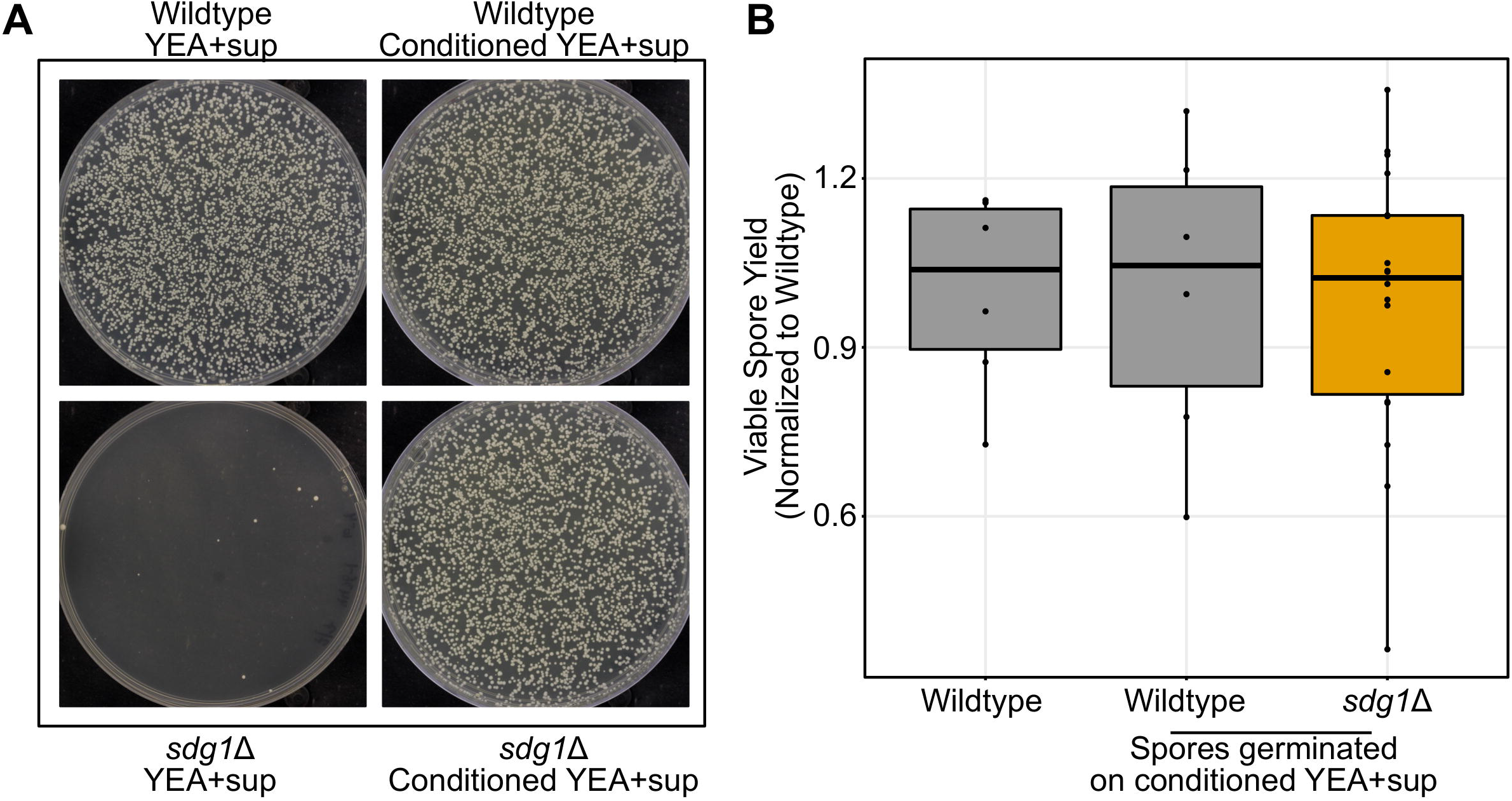
*sdg1*Δ mutants do not have a sexual reproduction defect if complemented by conditioned media. A) Spores from wildtype and *sdg1*Δ mutants were generated by incubating cells of each genotype on MEA plates for three days at 25°C. Equal dilutions of spores from wildtype and *sdg1*Δ mutants were plated to either YEA+sup or YEA+sup conditioned media and incubated for 5 days at 32°C. B) Viable spore yield assay showing on the y-axis the number of spores produced per yeast cell plated, normalized to the mean value for wildtype. Points display normalized results from a single replicate. The boxplot summarizes the underlying points and show first quartile, median, third quartile while the whiskers show the range of the data to a maximum of 1.5 times the interquartile range below and above the first and third quartile, respectively. Points outside the whiskers can be considered outliers. Cells were incubated on MEA plates for three days at 25°C in dense cell patches. Data shown for *sdg1*Δ combines three independent mutants performed on the same day. The wildtype on standard media was not different from wildtype on conditioned media (Mann-Whitney U test, p=0.937) nor were the *sdg1*Δ mutants (Mann-Whitney U test, p=0.923).

### Low density growth defect of *sdg1***Δ** mutants is not caused by auxotrophy

A similar quorum rescuable phenotype to that of *sdg1*Δ has previously been reported in *S. pombe* (Figure 7A). This phenotype is caused by the fact that some nitrogen sources (ammonium and glutamate) inhibit nutrient uptake from the environment in *S. pombe* [57]. As a result, auxotrophs may fail to import amino acids or nucleobases and thus fail to grow even when grown in media supplemented with the appropriate amino acid. However, this repression can be rescued by high cell density via oxylipins, which act as quorum molecules [58]. Because our mutants were constructed on a background auxotrophic for leucine and uracil, we hypothesized that *sdg1*Δ mutants may be a component of this previously described quorum phenotype. To test this idea, we analyzed the phenotype of *sdg1*Δ prototrophs and confirmed that mutation of *sdg1* did not itself confer auxotrophy (Figure 7 Supplement 1). To our surprise, prototrophic *sdg1* mutants still exhibited a strong low density growth defect (Figure 7B). As before, this low-density growth defect could be rescued by conditioned media (Figure 7C). Notably, the existing low density growth defect of *sdg1*Δ mutants was enhanced by addition of 5 g/L ammonium chloride in auxotrophs, but not in prototrophs (Figure 7 Supplement 2). Together, these data suggest that the quorum sensing pathway we have identified here is likely distinct from the previously described oxylipin pathway [58], although it does not rule out oxylipins as the quorum molecule driving this behavior.

**Figure 7.**
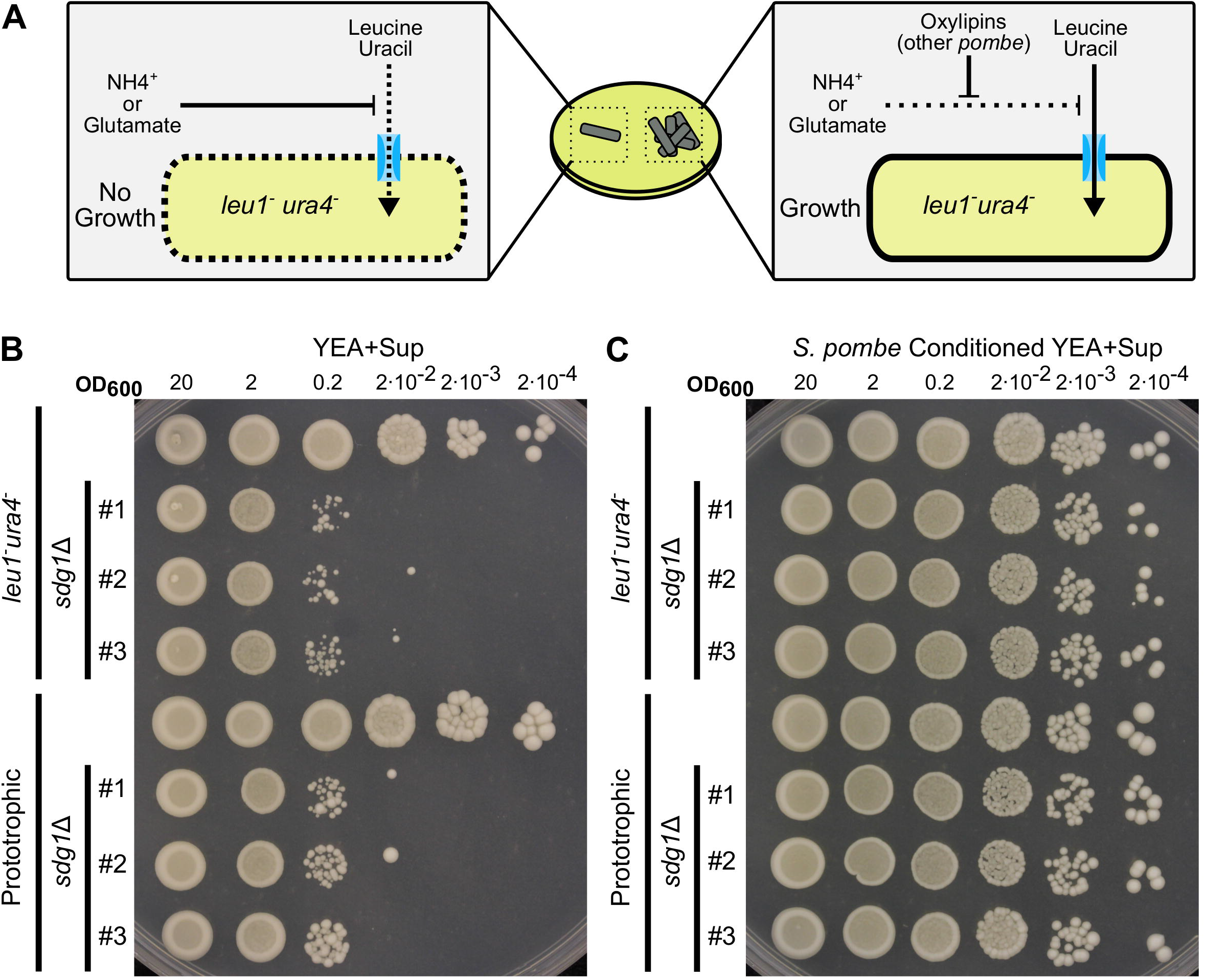
The *sdg1*Δ mutant phenotype is not the result of a previously described quorum effect. A) Diagram illustrating current model of oxylipin mediated quorum sensing in *S. pombe*. Some nutrient uptake is repressed by ammonia or glutamate in the media. As a result, auxotrophs may fall to grow even in the presence of the appropriate amino acids (left). However, this repression can be alleviated by oxylipins that are produced as a quorum molecule (right). Consequently, many *S. pombe* auxotrophs fail to grow at low cell density. B-C) Spot dilution assays with 5 μL spots plated. The initial leftmost spot is of OD_600_ = 20 culture and each successive spot is a 10-fold dilution, so that the final spot should be 10^5^ less concentrated than the first. Both experiments were conducted on the same day with the same dilution series of parent strain, three independent *sdg1*Δ mutants on the same background (*sdg1*Δ::kanMX4, *ura4*-D18, *leu1*-32), one wildtype strain and three independent prototrophic *sdg1*Δ mutants (*sdg1*Δ::kanMX4). Dilution series are from different cultures from those in Figure 5. B) Spotted to standard yeast extract agar (YEA+sup) and incubated for 4 days at 32°C. C) Spotted to conditioned yeast extract agar made from parent strain *S. pombe* (*ura4*-D18, *leu1*-32) and incubated for 4 days at 32°C.

### Shiny app allows navigation of sexual reproduction TN-seq data

As a resource for the *S. pombe* community, we have developed an interactive Shiny web app to navigate our sexual reproduction TN-seq data (Figure 2 Supplement 2). This is available at (https://simrcompbio.shinyapps.io/sex_tnseq_viewer/). Briefly, this app includes all data for coding genes that passed our site depth and gene insert number cutoffs. The figures produced are similar to those in Figure 2 but are automatically generated for a user selected gene. This app also integrates meiotic transcription data from Mata et al. [42]. This tool will allow the *S. pombe* community, and more broadly, the community interested in sexual reproduction to easily navigate this dataset.

## Discussion

Here we present a framework to allow direct experimental assessment of the genetic underpinnings of sexual reproduction in an organism in a single internally controlled experiment, without use of pre-existing resources like an arrayed deletion collection. Our assay identified 532 genes as important for sexual reproduction, including 217 without a relevant previously annotated phenotype in *S. pombe*. 164 of these novel candidate genes are conserved in vertebrates, suggesting that our approach could provide a powerful tool to identify unknown but conserved sexual reproduction genes and guide candidate-based approaches in systems where high-throughput approaches like TN-seq are not practical.

Like previous screens, there are both advantages and drawbacks to the TN-seq approach. Our assay was able to test a similar number of genes to those that have employed the *S. pombe* deletion collection (3418 genes here vs 3285 genes [14]). The set was not completely overlapping, in part because our statistical approach had limited power for shorter genes with small numbers of unique inserts. One strength of this approach is that largely neutral intergenic sites serve as a built-in control for wildtype sexual fitness. This is particularly important in *S. pombe* because we know from past work that wildtype meiosis is already surprisingly error-prone [52]. In addition, while our statistical approach in this study utilized the existing *S. pombe* annotation, the transposon insertion itself is annotation independent. Thus, future work exploring the subset of noncoding inserts with strong fitness defects may illuminate the contribution of noncoding regions of the genome to sexual reproduction in a way that would be impossible with a deletion collection approach. In fact, while our study did not systematically explore noncoding inserts, inserts into the noncoding RNA *sme2*, which is involved in pairing of homologs during meiosis [59], were also underrepresented after sexual reproduction (Mann-Whitney U test, p= 1.945 * 10^−12^) (Figure 2 Supplement 3).

One additional strength of the TN-seq approach is that even in species with existing deletion collections like *S. pombe*, secondary screens can be performed that would otherwise be impractical. For example, to further categorize the role of hits from our TN-seq screen, we could construct Hermes-insert libraries in a *rec12*Δ mutant background and repeat our screen. Genes that are required to repair meiotic double-strand breaks should no longer have a phenotype in a *rec12*Δ mutant, implicating them in the repair of DSBs [60]. In contrast, some genes may be synthetically required for meiosis in the absence of *rec12*, suggesting a role in *rec12*-independent chromosome segregation [61]. This process could be repeated for mutants with defined phenotypes throughout meiosis to generate a systematic genetic map of meiosis in *S. pombe*. In contrast, with traditional methods this would require reconstructing the full deletion collection on multiple genetic backgrounds, some of which already have low mating efficiency and thus would be very difficult to construct via mating. Further, TN-seq could be employed to model human infertility alleles in their orthologs in fission yeast and reassay the network of genes required for fertility. This would allow powerful interrogation of allele-specific gene network interactions for human-relevant conditions. TN-seq opens up new experimental avenues even in traditional model organisms that have the potential to provide powerful new insights into previously studied pathways.

### Comparison with past screens

Our screen varied in approach from two previous screens attempting to identify genes required for sex and meiosis, both of which used the *S. pombe* deletion collection. The first screen utilized iodine staining to specifically search for mutants that affected spore formation [36]. They identified 34 total genes spread over four categories, including genes required for zygote formation, iodine-reactivity, entry into meiosis, and for forespore membrane formation. Our screen assayed 29 of these genes and 17 were hits in our screen as well (58.6%). Ucisik-Akkaya et al. also identified a gold standard list of 13 known sporulation genes in *S. pombe* that had been experimentally tested. Their screen recovered 10 of the 11 they tested, while ours found 11 of the 12 we tested. Interestingly, we also identified a mutant that produced inviable spores that nonetheless stained with iodine. This suggests that iodine staining screens can miss genes that make inviable spores. This observation is not surprising, given that this has previously been observed for mutants that produce a limited number of inviable spores, such as *meu10* [62].

In contrast, Blyth et al. cytologically examined meiotic chromosome segregation and sporulation via microscopy [14]. They scored two criteria: percentage of yeast cells that went on to produce spores (% sporulation) and percentage of tetrads that formed four spores, each with a single chromosome II lacI-GFP dot and defined mutants as <20% sporulation (n=274) and/or <90% four spore tetrads (n=253). We recover a significantly larger proportion of Blyth et al.’s identified sporulation mutants (27.7%) than we do segregation mutants (15%) (Fisher Exact test, p=0.0005). This discrepancy is likely at least in part because our assay looks at the endpoint of meiosis: the production and fitness of spores. Because *S. pombe* has only 3 chromosomes, a simple model of random chromosome segregation predicts that spore viability remains relatively high [61]. At Blyth et al.’s 90% cutoff, spore viability is still predicted to be greater than 90%. This level of segregation defect will result in only a relatively subtle effect on fitness and thus on observed frequencies via TN-seq. Similarly, spore fitness can be affected by many processes other than failure to sporulate or chromosome segregation-*plb1* and *alg9* were both important, for growth post-sporulation for example.

We also note that our statistical approach was highly conservative. *atg11* was a borderline non-hit in our screen but proved to have subtle sexual reproduction defects when we examined it carefully. This suggests that our screen may have limited power to detect particularly subtle phenotypes and that there are likely additional components of the sexual reproduction machinery left to be identified in *S. pombe*.

### TN-seq can be used to search for density dependent growth mutants

In this study, we identified a gene (*sdg1*) that was previously annotated as essential but scored as both non-essential and involved in sexual reproduction in our screen. Through analysis of deletion mutants, we found *sdg1* to be non-essential, but required for growth at low density. This phenotype likely explains why this gene was originally misidentified as essential. Traditionally, tests of gene essentiality in fungi involve growth at low density. Tetrads from a heterozygote are dissected and a ratio of 2 live to 2 dead spores, where the live spores are wildtype, is used to infer that the gene has an essential function. However, this test requires the ability to grow at low density from a single spore to produce a visible colony. Therefore, mutants like *sdg1*Δ that are unable to grow at low cell density may incorrectly appear essential in this test. Indeed, when plated at low density, *sdg1*Δ spores failed to grow the vast majority of the time (Figure 6A).

More generally, most tests of spore viability also employ either tetrad dissection or random spore analysis. Both tests require spores isolated on plates to germinate and form countable colonies. A mutant with a defect in low density growth would fail to successfully form a colony and would be mistakenly counted as an inviable spore. Thus, our results suggest that it is possible that a subset of other mutants annotated with spore viability defects may be mischaracterized low-density growth mutants.

Further, this result, in conjunction with earlier studies of auxotrophy, suggests that *S. pombe* has a density-dependent growth program [57,58]. Auxotrophs fail to grow at low density in the presence of ammonium or glutamate but grow at high density. Cells lacking *sdg1* fail to grow at low density but grow well at high density or in conditioned media. While the defects caused by each appear independent, it remains unclear whether rescue of low-density growth is occurring through the same pathway in both scenarios. Future experiments can test whether there is one high density growth plan or multiple.

The *sdg1*Δ phenotype also suggests another use of our TN-seq assay to directly screen for mutants with low density growth defects. In our assay, our library of transposon-containing cells is plated at very low density so that they must form a microcolony in order to mate. Mutants that are capable of growth in dense culture, but are either incapable of forming a microcolony, or form smaller microcolonies, will fail to pass through our sexual reproduction assay. This is technically a false positive for our sexual reproduction assay, but a similar approach could be used specifically to screen for genes involved in low density growth. This low-tech sparse plating approach may achieve the same goals as the far more complex Droplet Tn-seq microfluidics and single cell amplification method recently described [63]. It is unlikely that mutants similar to *sdg1*Δ represent a large subset of our sexual reproduction hits, as only three other genes identified as essential were present as hits in our screen.

### Sexual reproduction TN-seq can identify mutants with impaired gamete health

Because our assay encompassed the entire process of sexual reproduction from mating all the way to spore germination, it enabled us to spot a new phenotype. Typically, spore germination is measured by assessing the fraction of spores that grow into colonies. This reduces spore health to a binary: spores either germinate or they do not. In contrast, our *plb1*Δ and *atg11*Δ mutants displayed a spore-specific growth defect where they germinated slowly but grew normally otherwise. Similarly, *alg9*Δ mutants also displayed a defect in spore growth, although they had a disadvantage growing vegetatively as well. Thus, “spore health” occurs as a range of phenotypes rather than simply a binary of alive or dead.

More broadly, spore health can potentially model gamete health in mammals, as both spores and eggs/sperm are products of meiosis. Declining gamete health is one of the main drivers and most poorly understood aspects of age-associated human infertility [64]. While we have no evidence that *plb1*Δ or *alg9*Δ mutant spores decline in health over time, the underlying challenges in building a gamete that is stable for long periods of time may be similar between fungal spores and mammalian eggs. One primary difference is that mammalian oocytes have not yet completed meiosis and chromosome segregation, while fungal spores have. Thus, this screen may be more likely to identify components of chromosome segregation-independent mechanisms that are specifically required to recover from meiosis and return to active mitotic growth. Future experiments utilizing TN-seq to better understand these mechanisms in fission yeast may enable us to eventually translate this understanding into mammalian systems and provide insight into the phenomenon of gamete health and human infertility.

### Exploring other members of the *Schizosaccharomycetes* clade and related species

We anticipate that this approach will allow highly productive future exploration of the evolution of sexual reproduction in the *Schizosaccharomycetes* lineage. Exploration of this complex could provide a model to study the evolution of the genetics of sexual reproduction over time, which is particularly interesting given the presence in *S. pombe* of a family of meiotic drivers called *wtf* genes [51,65,66] that appear to generate a selective pressure for less efficient meiosis [52].

In addition, a pan-genus understanding of sexual reproduction and meiosis in the *Schizosaccharomycetes* could provide valuable insight into the biology of *Pneumocystis jirovecii,* a fungal pathogen of humans that is a sister to the *Schizosaccharomycetes* lineage. *P. jirovecii* is responsible for 400,000 annual life-threatening infections, typically in conjunction with HIV/AIDS [67]. Because *P. jirovecii* cannot be grown as a free-living organism, studies in related organisms will be critical to understand its biology. This is particularly relevant as recent studies have suggested that *P. jirovecii* may be undergoing sexual reproduction inside a human host in order to produce sexual spores that can be transmitted to a new host to establish infection [68–70]. Therefore, understanding spore formation and germination, as well as developing inhibitors for it, could be critically useful in blunting *P. jirovecii* infection. More broadly, spores often serve as infectious particles for many fungal pathogens [71] and thus understanding the genetic network required to produce functional spores could be helpful in preventing infection.

## Material and methods

### Data availability

All original data is available at the Stowers original data repository. Next-gen sequence data is available at the SRA under accession number PRJNA758956.

### Strain construction

We transformed all *S. pombe* strains using a standard lithium acetate transformation protocol [72]. We constructed all mutants with at least two independent transformations using independent starting cultures and we then PCR validated to confirm correct insertion of the deletion cassette and loss of the original coding sequence. Strains are listed in Table S2. We cryopreserved all strains at −80°C in 20% glycerol and revived them on yeast extract agar (YEA+SUP) at 32°C prior to conducting experiments. For deletion of *ifs1*Δ, *atg11*Δ, *plb1*Δ, and *alg9*Δ, we amplified the kanMX4 allele amplified from the *S. pombe* deletion collection strain with approximately 500 base pairs of homology up and downstream. Primers for each can be found in Table S3. For deletion of *sdg1*Δ and *ggt1*Δ, we constructed deletion cassettes via overlap PCR with kanMX4 from plasmid pFA6 [73]. Primers for these overlap PCR reactions are also listed in Table S3.

We constructed prototrophic *sdg*1Δ mutants by crossing our auxotrophic *sdg1*Δ mutants to strain SZY513 and collecting the spores. We then plated and grew them on YEA+SUP+G418 to select for *sdg1*Δ::kanMX4 and then replica plated to minimal media to select for prototrophs (for SZY4831). Alternatively, we initially grew them on minimal media and then replica plated to YEA+SUP+G418 (for SZY4834 and SZY4837).

To generate our *ifs1*Δ strain in an Isp3-GFP background (SZY5082), we transformed the *ifs1*::kanMX4 deletion cassette into SZY4975 (*h*^90^ *isp3*-GFP *leu1*::mCherry-psy1: FY39320 from the Yeast Genetic Resource Center) [74].

### TN-seq Library Preparation

We prepared transposon insertion libraries using a modified method from the one presented in Guo et al [28]. We transformed plasmid pHL2577 carrying the Hermes transposon and pHL2574 carrying the NMT-transposase into SZY643 (*h^90^*, *leu1*^−^, *ura4*^−^) and selected transformants on EMM+ade+his+lys+Thiamine media. Rather than picking individual colonies, we scraped the transformation plate and collected all transformants to inoculate a 50 mL EMM+ade+his+lys+Thiamine culture to grow overnight until saturated. We then washed 3 times with EMM+ade+his+lys (lacking thiamine) to remove any residual thiamine and resuspended in 25 mL of EMM+ade+his+lys. We used 500 µL to inoculate 50mL of EMM+ade+his+lys to induce transposition. We then serial cultured over 4 days by adding 1 mL (day 2) or 1.5 mL (days 3 and 4) to 50 mL of fresh EMM+ade+his+lys. We selected for cells with transposition events by first spinning down 10 ml of this final culture (3000 rpm, 5 min), removing 9 mL of the supernatant and resuspending the cells in the remaining 1 mL of supernatant. We used this to inoculate 100 mL of YEL+Sup+5-FOA and grew overnight to counter-select against the initial Hermes plasmid. We then spun 50 mL down as previously described and resuspended the cells in 5 mL of the supernatant. We inoculated this into 500 mL of YEL+Sup+5-FOA+G418 and grew it for 2 days until saturation. The resulting *ura*^+^ and G418^R^ cells should have undergone transposition but lost the pHL2577 plasmid. We collected cells and split them to freeze in 20% glycerol as well as to prepare DNA.

We prepared DNA using a Qiagen Genomic Tip column with the standard yeast protocol except we extended the lyticase and proteinase K treatment to 24 hours on a shaking incubator. We then digested the purified DNA using either MseI (10,000 U/mL) or a combination of HpaII (10,000 U/mL), TaqI-V2 (20,000 U/mL), and HpyCH4IV (10,000 U/mL). Our digests used 10 µg of DNA in 500 µL of volume overnight at 37°C in CutSmart buffer with 15 µL of each enzyme, except that TaqI-V2 was not added until after the overnight digest, after which we raised the temperature to 65°C for 4 additional hours. We cleaned and size-selected the digested DNA using SPRIselect beads by washing with 0.5 volumes of SPRIselect beads, pelleting on a magnet and retaining the supernatant, and then adding 0.2 volumes of beads and precipitating on beads again. Finally, we washed and eluted the DNA from the beads with 500 µL of water.

We then end-ligated on linkers that contained unique, random barcodes using a reaction containing 490 µL of the cleaned, size-selected DNA, 88 µL of 10x ligation buffer, 143.5 µL of water, 153.5 µL annealed barcoded linker, and 5 µL T4 DNA ligase (400,000 U/mL). We then divided the reaction evenly between 32 PCR tubes and incubated for 16 hours at 16°C. We prepared the barcoded adapter as in Guo et al. [28] using oligo oSZ2760 that contains random nucleotides for the unique barcode, and either oSZ2485 for the MseI digested DNA or oSZ2499 for the combination HpaII, TaqI-V2, and HpyCH4IV digested DNA (Supplemental Table 3). We annealed the oligos together by mixing at a concentration of 10 µM each in 1x HF PCR buffer and denatured at 95°C for 1 minutes, followed by decreasing the temperature 10°C for 7 minutes, and then further decreasing the temperature by 10°C every 7 minutes until it reached 20°C. We stored the annealed adapter at −20°C. The unique barcodes allow us to directly count the number of inserts present in our library after sequencing [30].

At this stage, we split the DNA from each digest into two pools (four pools total). We amplified these pools via PCR with Phusion polymerase and 15 different Hermes-specific oligos (oSZ2104-2118, Supplemental Table 3) with Illumina adapter tails. We used 3 reactions per oligo per pool, resulting in a total of 90 PCR reactions for each enzyme mix. In addition, we ran 3 linker-only controls. Each PCR reaction contained 24.5 µL water, 10 µL 5x HF buffer, 1 µL dNTPs (10 mM), 8 µL linker ligated DNA, 1 µL linker oligo oSZ2483 (10 µM), 5 µL Hermes-specific oligo (2 µM) or water, and 0.5 µL Phusion polymerase (2,000 U/mL). We amplified inserts using the same thermocycler settings as Guo et al. [28]: 94°C for 1 min, 6x (94°C for 15 sec, 65°C for 30 sec, 72°C for 30 sec), 24x (94°C for 15 sec, 60°C for 30 sec, 72°C for 30 sec), 72°C for 10 min, 4°C hold. For each primer pair and pool, we then combined the three reactions to form subpools, a subset of which we ran on a gel for validation. We then added the remaining Illumina adapters and barcodes with another round of PCR. This PCR reaction contained 28.5 µL water, 10 µL 5x HF buffer, 1 µL dNTPs (10 mM), 2.5 µL oSZ2128 (10 µM), 2.5µL barcode oligo (oSZ2129-2136, 2763-2766; 10µM), 5 µL of insert PCR pool, 0.5 µL Phusion polymerase (2,000 U/mL) and we performed it at 94°C for 2 min, 5x (94°C for 30 sec, 54°C for 30 sec, 72°C for 40 sec), 4°C hold. The individual pools are given a unique barcode (2 barcodes per enzyme digest mix and 4 total barcodes per DNA library). We then cleaned the DNA using 0.75 volumes of SPRI beads, precipitated on a magnet, washed, and eluted. We confirmed the DNA library by PCR using oligos to the ends of the Illumina adapters. This reaction consisted of 15.75 µL water, 5 µL 5x HF buffer, 0.5 µL dNTPs (10 mM), 1.25 µL oSZ2176 (10 µM), 1.25 µL oSZ2177 (10 µM), 1 µL library DNA, 0.25 µL Phusion polymerase (2,000 U/mL). We ran this PCR at 98°C for 30 sec, 35x (98°C for 10 sec, 62°C for 30 sec, 72°C for 40 sec), 72°C for 10 min, 4°C hold. Finally, we quantified this DNA library using a Qubit and determined fragment sizes using a Bioanalyzer. Sequencing was performed on a NextSeq instrument at the Stowers Molecular Biology Core. Raw sequencing reads are available at PRJNA758956 on the Sequence Read Archive.

### TN-seq Mating Assay

We revived a frozen aliquot (1.8 mL) of our *S. pombe* Hermes TN-seq library by thawing it on ice and growing it in 50 mL of saturated culture in YEL+sup overnight. We used part of this culture to prepare a second round of pre-sex sequencing libraries (see above), while we diluted the remainder 1000-fold for plating. We plated 250 μL of this diluted culture (an estimated 18,500 cells) to each of 144 large (150 mm) MEA plates. We then incubated these plates at 25°C for 9 days. After incubation, we collected cells from the MEA plates by physically scraping cells from all 144 plates. We specifically isolated spores using a standard glusulase prep and then allowed them to germinate in YEL+sup liquid culture for either a “short” (single 1:100 dilution of spores into rich media and grown for 24 hours) or “long” (1:1000 dilution of spores into rich media grown for 24 hours and diluted 1:100 into rich media and grown for another 24 hours) outgrowth period. We then collected cells and isolated DNA using the Qiagen Genomic Tip kits. This DNA was used to prepare sequencing libraries as described above.

### Data Analysis

We initially processed sequencing reads using a custom R script to identify reads with the correct linker sequence. We kept only reads with a perfect match to the linker sequence. We saved and linked the unique barcode sequence added with the linker to the read while the entire linker sequence was trimmed off. We then imported reads with the proper linker sequence into Geneious Prime (v. 2020.2). We removed reads if they did not contain sequence matching perfectly to the entire end of the Hermes transposon from oligo to insertion site, 61-85 bp depending on the oligo. We did this using the “Separate Reads by Barcode” function in Geneious which also trims the Hermes sequence from the read at the same time. We mapped the remaining reads to the *S. pombe* genome (version ASM294v2) using the Geneious aligner within Geneious. Non-unique mappings were not allowed. We then output this alignment and ran it through a custom R script to generate a list of read start positions (i.e. Hermes insert sites) with a corresponding depth of unique barcodes at that position for each sequencing library. We used custom Perl scripts to assign genes to each insert site based on the *S. pombe* annotation (version ASM294v2), to combine depth files from the 4 individual sequencing runs for each treatment, and to track positions where inserts were present in the initial library but missing from subsequent libraries. We then used R to perform statistical tests. To limit low-frequency sampling issues, we dropped any site with eight or less unique inserts in our pre-sex dataset. We calculated a log_10_ fold change for post-sexual reproduction over pre-sexual reproduction frequencies. To perform this analysis, we set all 0 insert sites in the post-sex libraries to 1. This artificially reduces the magnitude of phenotypes observed but allows log adjustment of our data. We then filtered out all genes with less than 5 insert sites. We used a Mann-Whitney U test to compare the distribution of fold-changes for the remaining genes to that of the inserts annotated as intergenic. Finally, we performed multiple test correction using a Bonferroni correction.

### Viable Spore Yield

We started overnight liquid cultures from a single colony in 5 mL of YEL. The next day, we spotted 50 μL saturated culture in a single patch (for dense patches) or spread 111 μL of a 10^−3^ dilution onto an entire SPAS or MEA plate (for sparse patches). The sparse dilution is scaled to reproduce the same plating density used in the initial TN-seq assay but on a smaller petri plate. At the same time, we made serial dilutions of these cultures and plated to YEA+sup plates to determine the number of colonies per mL and thus the number of colony forming units we had spotted on the mating plates. We incubated the mating patches at 25°C. After either 3 or 9 days, we scraped the entire patches off the plates and used a standard glusalase prep to isolate spores [48]. We then performed a serial dilution and plated spores to YEA+sup plates in order to count the number of spores present in that patch. We calculated viable spore yield by dividing the number of viable spores present in the patch by the number of yeast cells originally plated. Additionally, we have normalized each experiment to the mean VSY of a control (SZY643) experiment done at the same time.

### Iodine Staining

We spotted 100 μL of culture from saturated overnight liquid cultures onto MEA plates and incubated them for 3 days at 25°C. We then inverted these plates over iodine crystals in a chemical fume hood until positive control cell patches had stained. We scored and imaged these plates immediately.

### Aneuploidy Assay

We generated test strains to measure rate of aneuploidy for chromosome 3 by crossing our *plb1*Δ and *alg9*Δ mutants to a strain (SZY2465, *ade6*Δ::hphMX6, *h*^−^) carrying *ade6* deleted with hphMX6. We grew overnight cultures of each strain in liquid YEL+sup media at 32°C. We then mixed equal volumes of each parent strain and spotted them to MEA. We incubated these plates for 3 days at 25°C. We then collected spores, glusulase treated, and plated spores to YEA+sup+Hyg plates. We picked and master plated spore colonies to YEA+sup plates. We then replica plated to identify progeny that were *ade6*Δ::hphMX6, *ura4^−^,h^−^*. We also generated a second test strain by crossing our *plb1*Δ and *alg9*Δ mutants to an *h*+ strain SZY128 (*ade6^−^, leu1^−^, his5^−^, h^+^*). We carried this assay out as described above except that spores were plated to YEA+sup rather than YEA+sup+Hyg. From this cross, we selected progeny that were *leu1^−^, his5^−^, G418^R^,h^+^*. From each cross, we also selected a strain with the same genotype but with a wildtype allele of *plb1* and *alg9*.

To assay aneuploidy, we crossed our *h*^+^ and *h*^−^ mutant strains to each other and our *h*^+^ and *h*^−^ wildtype strains to each other on MEA for 3 days at 25°C. We collected spores, glusulase treated, diluted them, and plated on YEA+sup plates. We picked entire spore colonies to make a YEA+sup masterplate after 5 days. This plate was replica-plated to each test media and chromosome 3 disomes were scored as colonies that were both ade+ and hygromycin resistant. We used the remaining markers to verify that strains were mating as expected (approximately 50:50 inheritance of unlinked markers). We did not score crosses with obvious strong divergence from 50:50 inheritance of the unlinked markers.

### Competition Assay

To conduct spore competition assays, we mixed approximately 100,000 viable parental spores (*h^90^*, *leu1*^−^, *ura4*^−^) with either 100,000 spores of our tester mutant strain or 100,000 spores of a presumably neutral *wtf12*Δ pseudogene deletion that was also G418-resistant. Spore numbers were quantified by plating cells on YEA+sup and determining the colony forming units. We then diluted the spore mix into 5 mL of YEL+sup, took a sample and plated to determine the initial frequency. We replica plated these plates to YEA+sup+G418 to count the proportion of G418-resistant and susceptible colonies. We also grew the 5 mL mixed culture overnight with shaking. The next day, we diluted and plated these spores to YEA+sup to acquire approximately 200 colonies per plate. Once they had grown, we replica plated these plates as well to determine the final frequency of each strain in the mix.

For vegetative competition assays, we started 5 mL overnight cultures using single colonies of the parental strain, our test mutant, and the *wtf12*Δ deletion strain. The next day, we determined OD_600_ for these cultures using a spectrophotometer and diluted them back to an OD_600_ of 0.001. We then mixed each with an equal amount of adjusted parental strain in 5 mLs of fresh YEL+SUP. We sampled and plated this initial mix, as above, and then grew the competition cultures overnight with shaking. Finally, we plated and replica plated to determine the proportion of each strain found in the initial and final mix.

### *S. pombe* functional annotation

We acquired *S. pombe* functional annotations (FYPO) from PomBase on December 15, 2020 [41]. To compare to our TN-seq data, we selected only annotations representing deletion phenotypes.

### Media

We made YEA+sup plates with 5 g/L Bacto Yeast Extract, 30 g/L Dextrose, 20 g/L Bacto Agar, 200 mg/L Adenine, 200 mg/L Histidine, 200 mg/L Leucine, 200 mg/L Lysine, 200 mg/L Uracil. We made YEL+sup in exactly the same way but omitting the Bacto Agar.

We made *S. pombe* conditioned media by starting a 5 mL YEL+sup culture from a single colony of SZY643 (*h^90^*, *leu1*^−^, *ura4*^−^). We grew that culture overnight at 32°C and diluted it back into 50 mL of fresh YEL+sup. We then grew that culture overnight as well and then diluted it back into 500 mL of YEL+sup and grew it overnight again. We then collected that entire culture via centrifugation at 5000g for 10 minutes at 4°C. We filtered the supernatant through a 0.2 μm vacuum filter and stored it at 4°C until use. We then made conditioned media using the standard YEA+sup recipe, except that we replaced 50% of the liquid component with the filtered culture supernatant described above prior to autoclaving. We stored plates at 4°C in sealed plastic sleeves until use.

### Spot Dilution Assays

To perform spot dilution assays, we grew cultures overnight in YEL until they reached saturation. We then measured the OD_600_ using a spectrophotometer. We pelleted enough culture to produce an OD_600_ of 20 in a final concentration of 1 mL by spinning at 3000 rpm for 5 min. We then removed the supernatant by pipetting and resuspended in 1 mL of water to remove any effects of conditioning in the residual media. We then performed 10-fold serial dilutions to a final concentration of OD_600_ =2*10^−4^. We spotted 5 μL spots of each intermediate dilution onto the appropriate media and incubated plates at 32°C, or 25°C for MEA media, until we took pictures as detailed in individual figures.

### Microscopy

To perform live imaging, we spread isolated spores onto a YEA+sup plate using beads. Immediately after the plates dried, we removed the beads from the plate and removed a circular punch of agar from the plate using a 1271E Arch Punch as previously described [75]. We inverted the agar and placed it top side down into a 35 mm glass bottom dish (No. 1.5 MatTek Corporation) containing a damp kimwipe and sealed the lid with vacuum grease. We then mounted the dish in a stage top incubator (Oko Lab) to maintain a temperature of 32°C. We imaged continuously on a Nikon Ti2-E widefield microscope, taking brightfield exposures using a Plan Apochromat Lambda 60x objective (1.4 NA) every 10 minutes for either 24 or 48 hours total. We processed images with Fiji (https://imagej.net/software/fiji/) to remove drift using stackregj (when necessary) and cropped to smaller image sizes for convenience. We manually scored steps throughout spore germination and division using Fiji and by manually drawing ROIs.

To take still images of cells plated to MEA we scraped a small portion of cells from the plate, resuspended in lectin (1 mg/mL) and imaged on a slide. To image cells grown on SPA we used the punch method described above. Cells were imaged using an Axio Observer.Z1 (Zeiss) wide-field microscope with a 40x C-Apochromat (1.2 NA) water-immersion objective. To excite GFP, we used a 440-470 nm bandpass filter, reflected the beam off an FT 495 nm dichroic filter into the objective, and collected emission using a 525-550 nm bandpass filter. We collected emission onto a Hamamatsu ORCA Flash 4.0 using μ

### Deep Learning

We segmented images using the Mask R-CNN [76] convolutional neural network as implemented in the pytorch [77] model zoo. Mask R-CNN is a neural network pre-trained to identify and segment everyday objects from a large set of images. Since the Mask R-CNN network was not trained specifically on cells, we fine-tuned it to identify *S. pombe* cells and spores. We did this by manually outlining or filling spores and cells in 12 transmitted light images and then retraining the network. During training, we sliced images into random 400×400 patches and augmented this with rotation and mirroring every iteration. We ran training for 200 epochs. We performed inference by slicing an input image into 400×400 patches, running the network on each patch, and reassembling the predicted patches back to the same size as the original image. The training and inference packages were written in python using pytorch as the deep learning library. We performed computation on a workstation with Ubuntu Linux 18.04 and an NVIDIA Quadro RTX 8000 for a GPU.

We loaded the output (predictions) of the deep learning network into Fiji (https://imagej.net/software/fiji/) for analysis. First, we thresholded the predictions manually to create a binary mask. We then registered this mask for movement when necessary using “Stackregj.” Next, we turned this mask into a list of ROIs using “Analyze Particles.” We measured these ROIs in order and acquired all Fiji measurements. These measurements included, but were not limited to, the area, the aspect ratio, and the minimum and maximum axes of a fit ellipse, among others. We saved these measurements as a .csv file and loaded them into a jupyter notebook written in-house to track spores from one frame to another.

## Acknowledgements

We thank members of the Zanders lab for comments on the manuscript. We thank the Levin lab for providing Hermes plasmids as well as advice on TN-seq library preparation. This work was funded by the Stowers Institute for Medical Research, a Searle Scholars Award to SEZ, as well as K99/R00 funding to SEZ (R00GM114436) and DP2 funding to SEZ (DP2GM132936) from the NIH. Strain FY39320 was provided by the NBRP (YGRC).

---

**Figure 2 Supplement 1.**
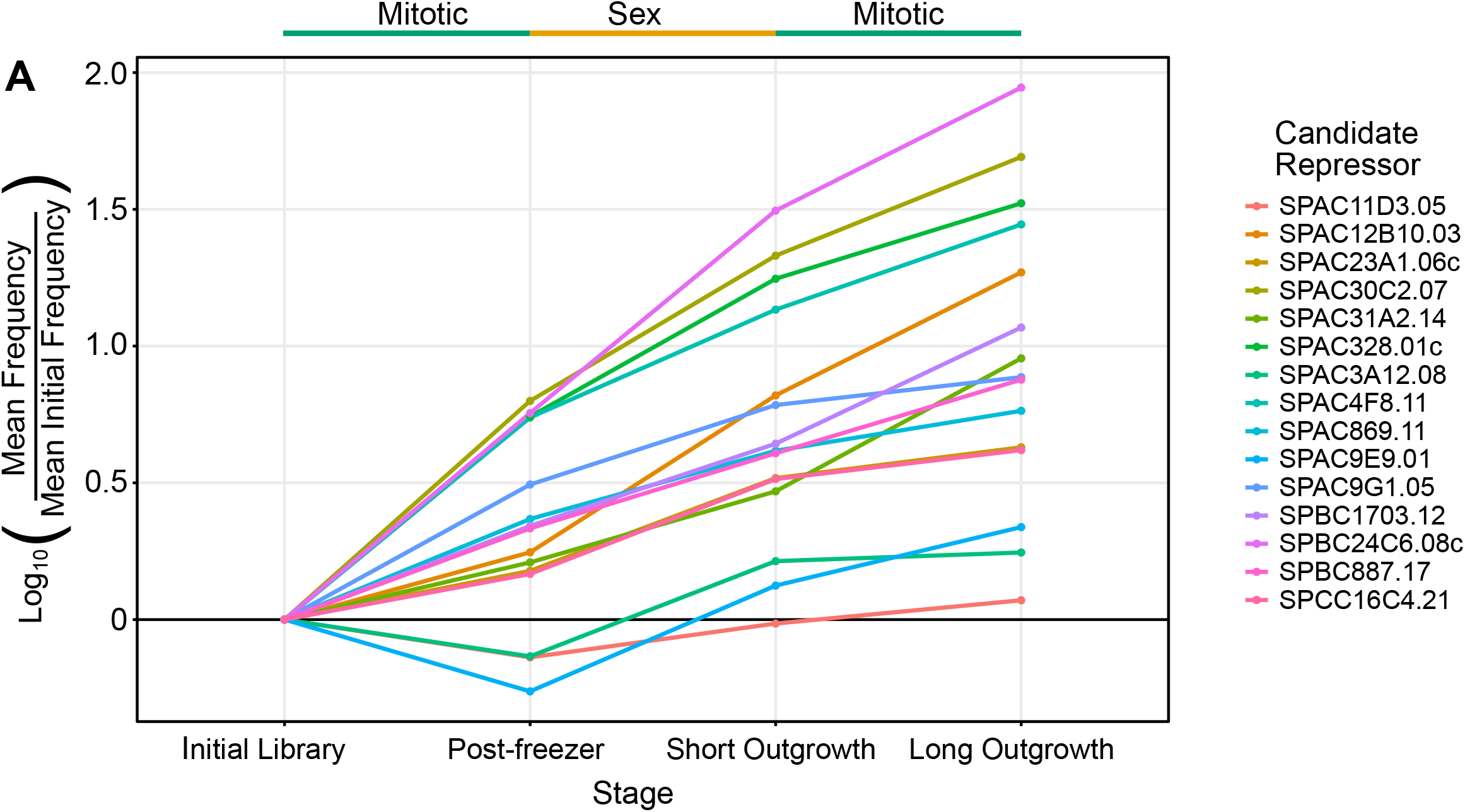
Candidate suppressors largely appear to have growth rate advantages. TN-seq insert frequencies for 15 genes identified as candidate repressors of sexual reproduction. The y-axis displays the log_10_ adjusted ratio of mean insert frequency across a gene to the mean insert frequency for that gene at the first sequencing step. The first and third segments are vegetative growth, while the second is a sexual reproduction step.

**Figure 2 Supplement 2.**
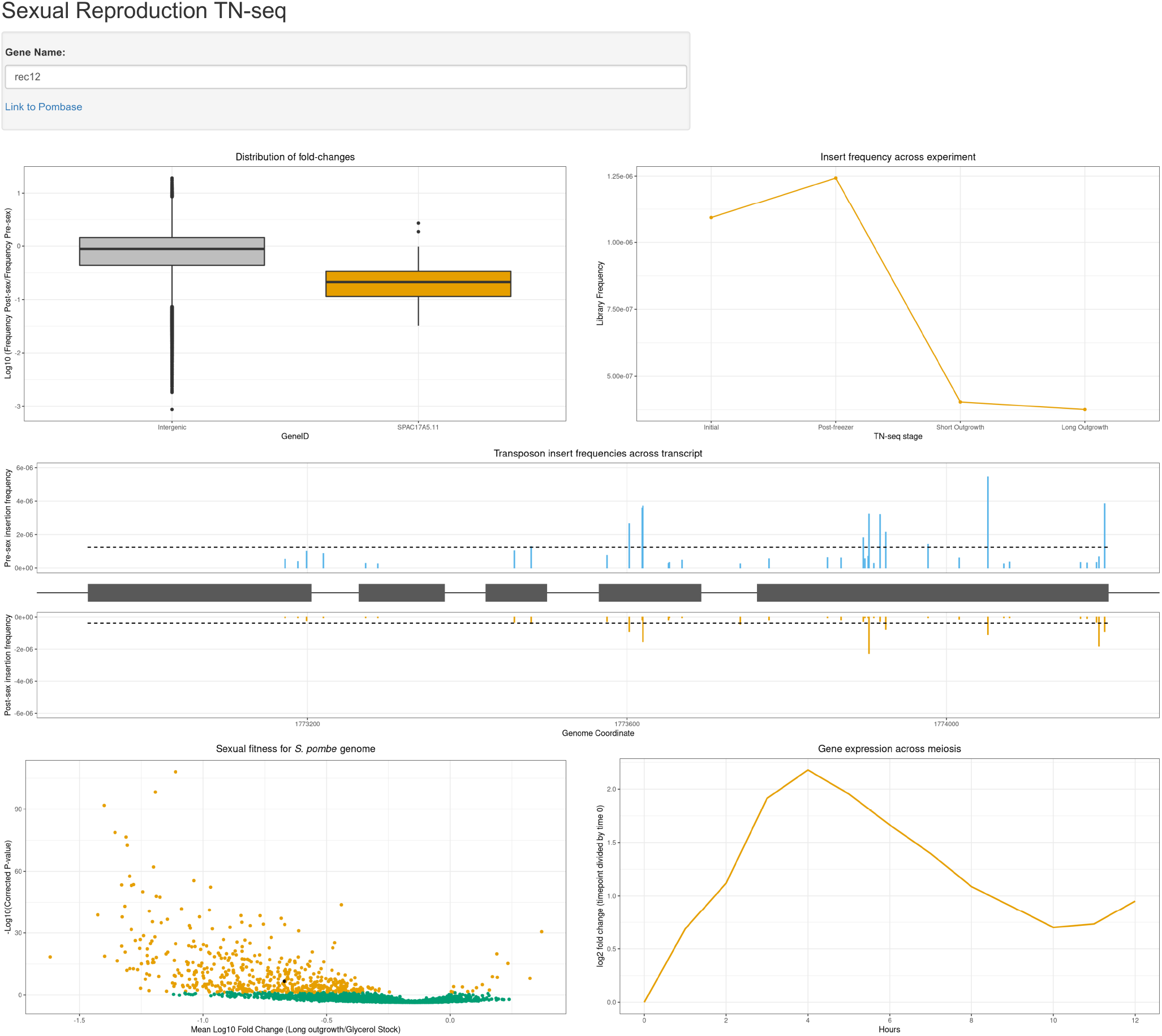
Sexual reproduction TN-seq data can be visualized using a publicly available Shiny app. Screenshot of a publicly available interactive Shiny app that visualizes data from the TN-seq screen. There are five plots, displaying the distribution of insert fold changes within a gene (as in Figure 2B), the mean insert frequency within a gene over time (as in Figure 2 Supplement 1), the site-by-site frequency across a gene (as in Figure 2A), a volcano plot summarizing the entire experiment (as in Figure 2C), and a plot of transcription throughout meiosis for that gene from the Mata et al. dataset [42] (as in Figure 2D). This app will accept *S. pombe* gene names, either as systematic names or common names.

**Figure 2 Supplement 3.**
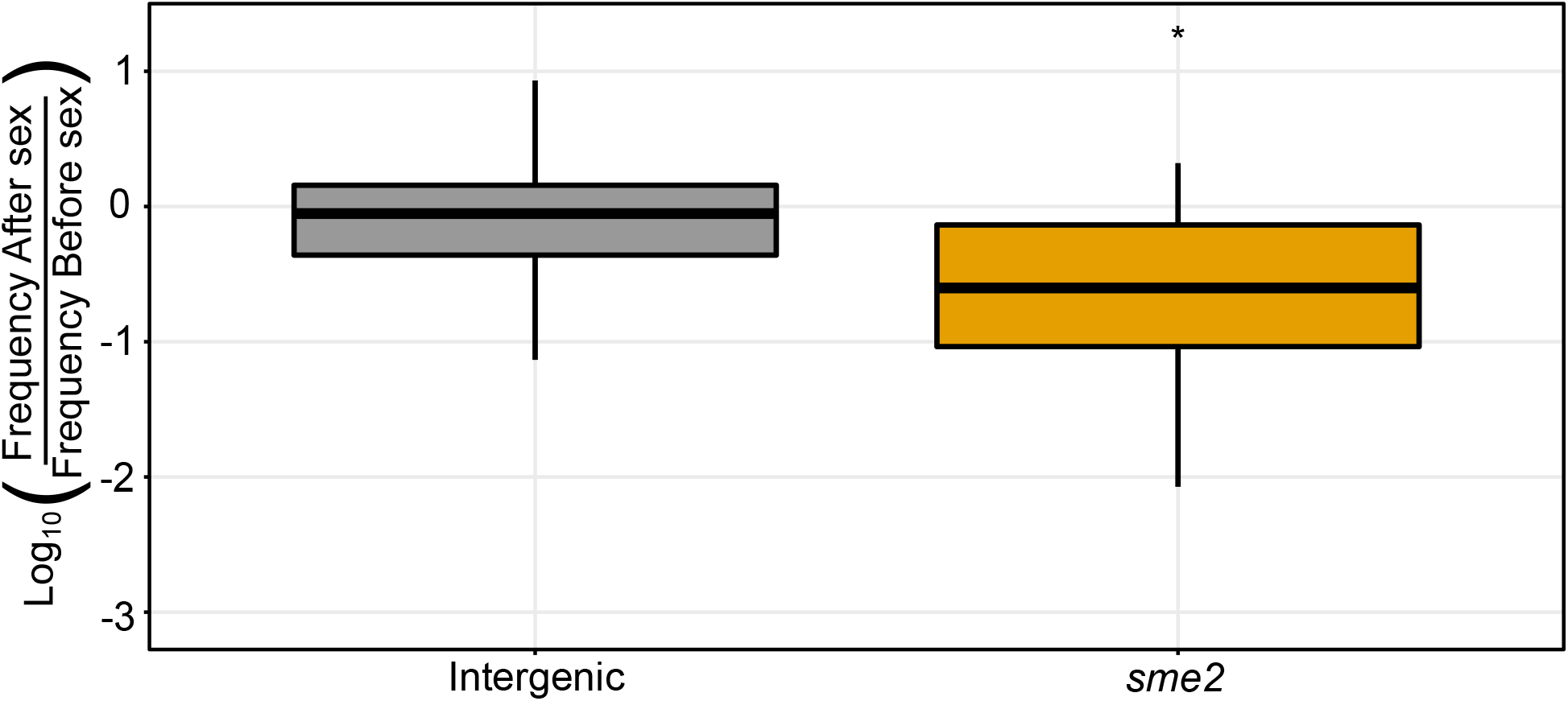
Inserts into the noncoding RNA *sme2* are significantly underrepresented after sexual reproduction. Boxplot displaying distribution of log_10_-adjusted fold changes in insert density after sex (ie. frequency after sex/frequency before sex). Boxplots show first quartile, median, third quartile. The whiskers show the range to a maximum of 1.5 times the interquartile range above and below the first and third quartile, respectively. Outlier data points (outside the whiskers) are not displayed. This results in the removal of 5,716 of 235,578 intergenic sites, and 0 of 78 from *sme2*. Inserts in intergenic regions are indicated in grey and inserts into the known meiotic noncoding RNA *sme2* are shown in orange.

**Figure 3 Supplement 1.**
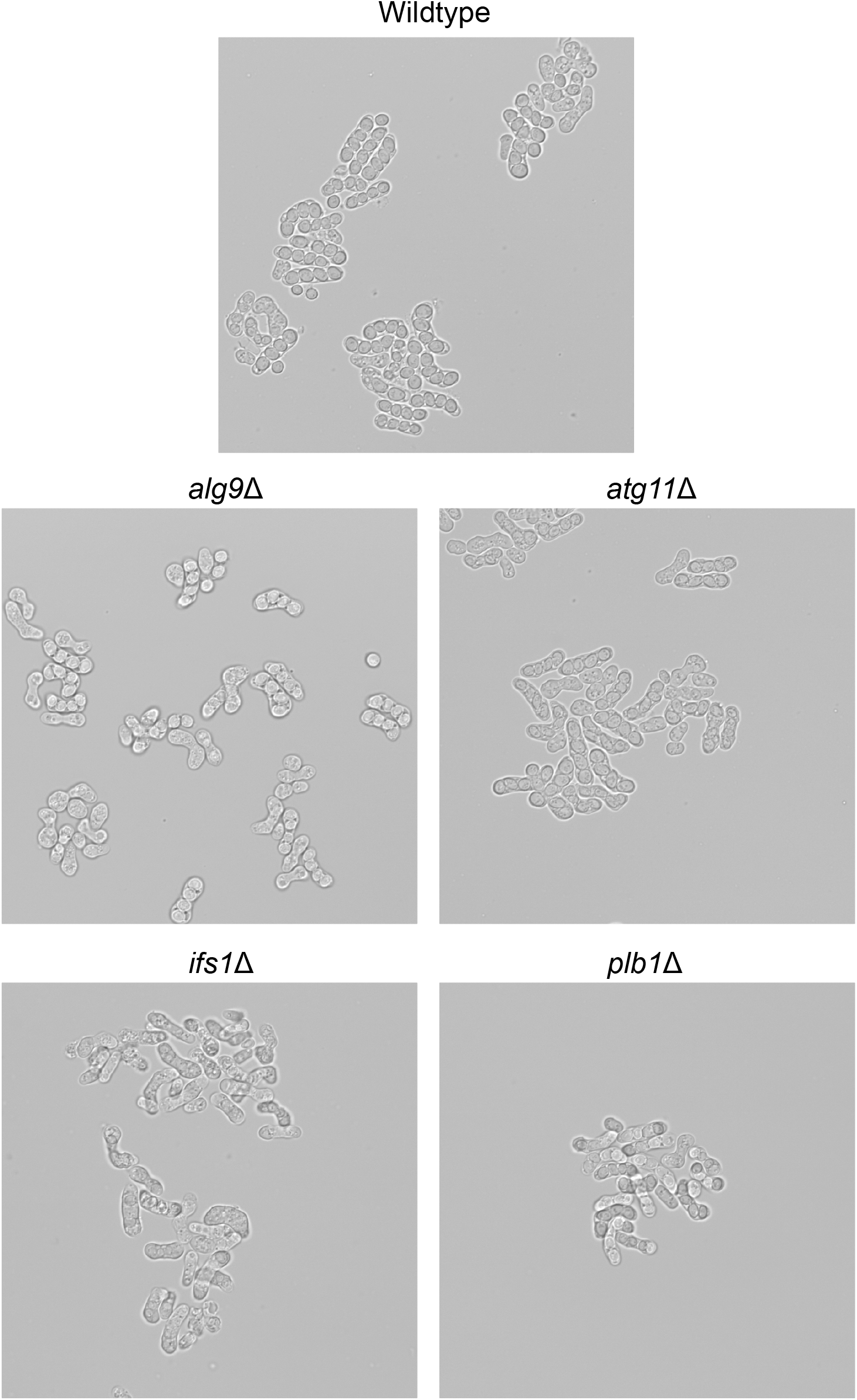
Candidate mutants produce sexual spores. Imaging on an AXIO Observer.Z1 (Zeiss) wide-field microscope with a 40x C-Apochromat (1.2 NA) water-immersion objective after of wildtype and mutant *S. pombe* sexual spores/asci produced after 2 days at 25°C on MEA.

**Figure 3 Supplement 2.**
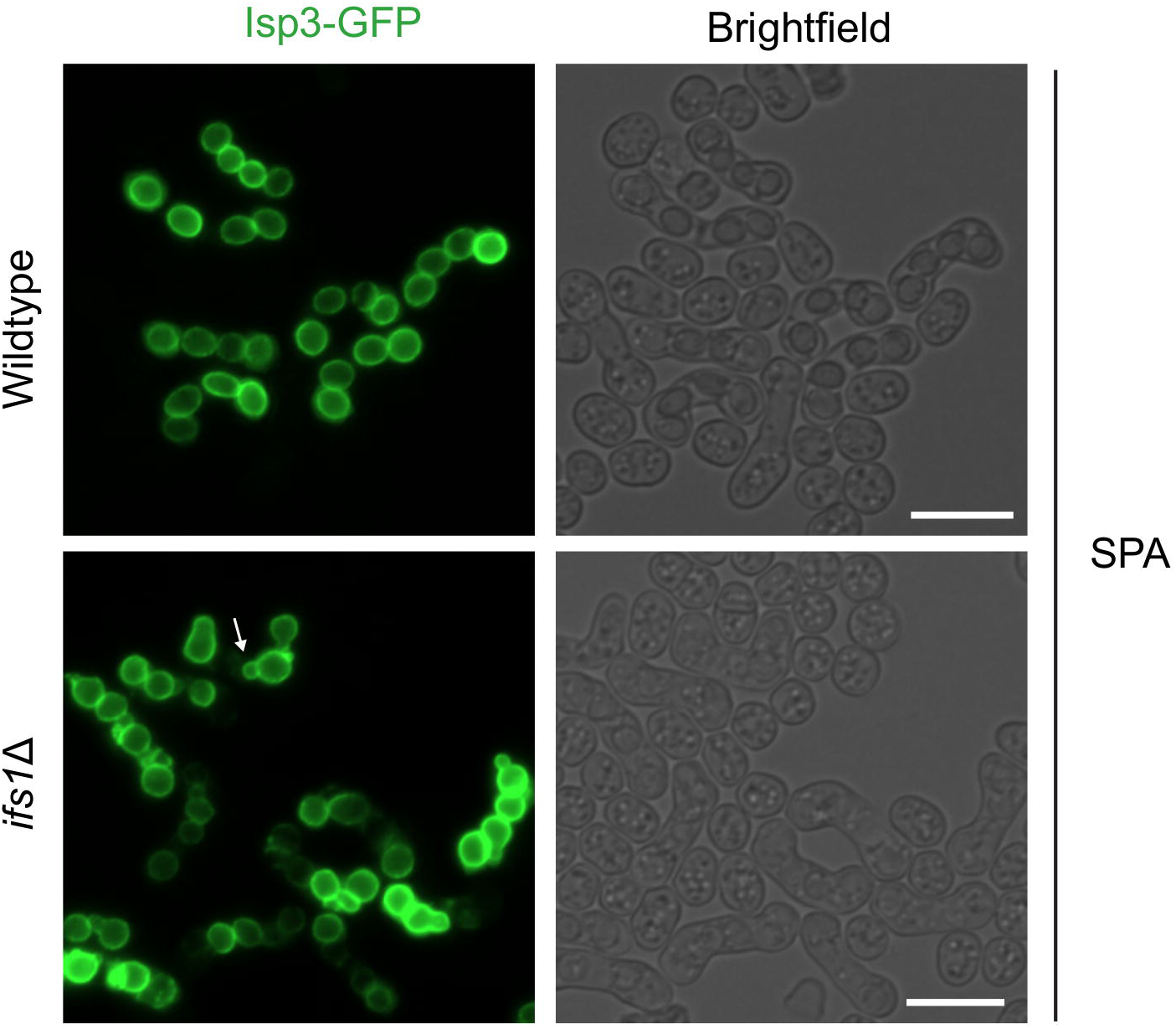
*ifs1*Δ spores are irregular and snowman-shaped in SPA media. Isp3-GFP was visualized in wildtype and *ifs1*Δ mutants on an AXIO Observer.Z1 (Zeiss) wide-field microscope with a 40x C-Apochromat (1.2 NA) water-immersion objective after incubation for 2 days at 25°C on SPA media. Scale bars indicate 10 microns. Arrow indicates “snowman” spore.

**Figure 3 Supplement 3.**
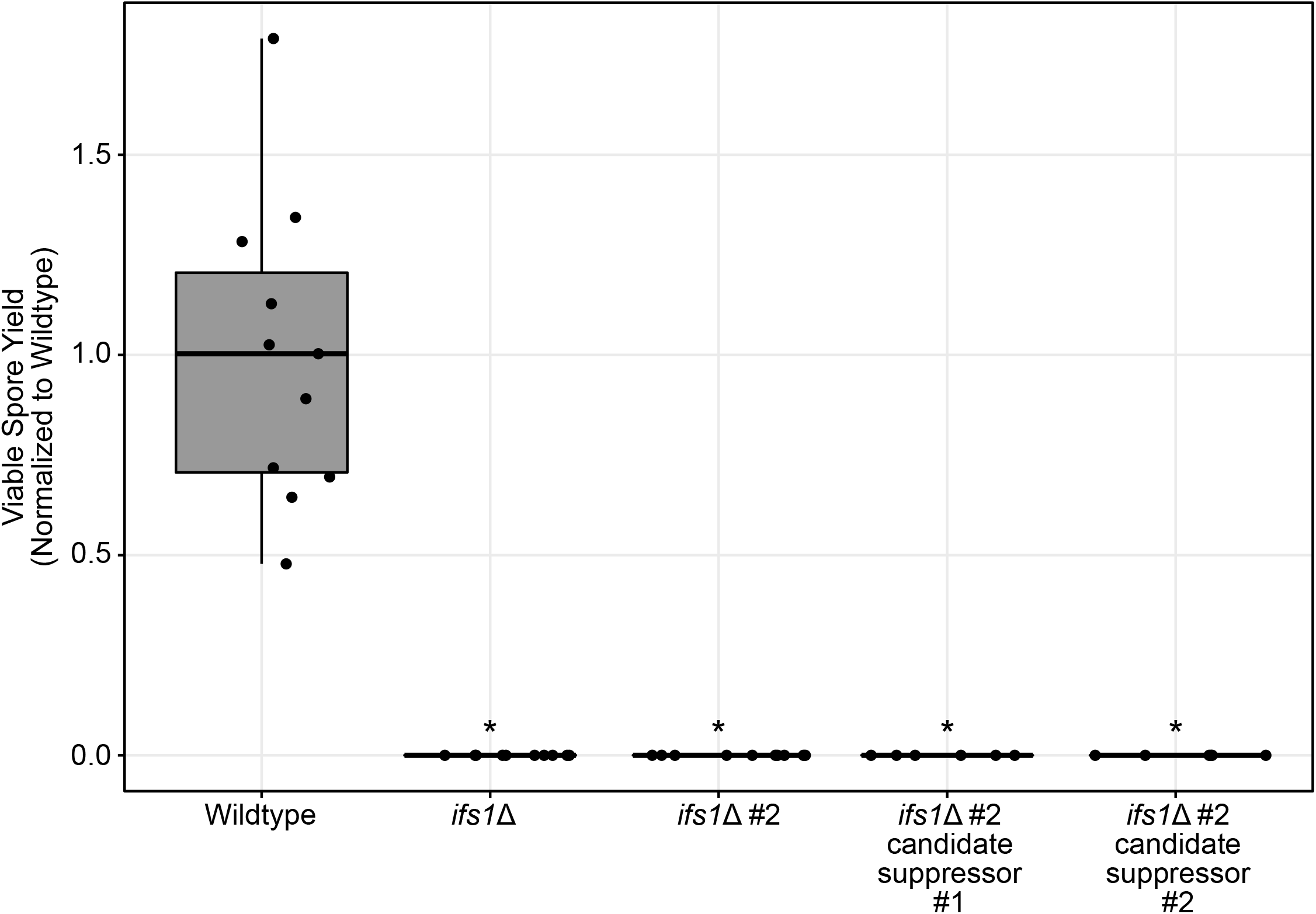
Surviving *ifs1*Δ spores retain their mutant phenotype. Viable spore yield assay showing on the y-axis the number of spores produced per yeast cell plated, normalized to the mean value for wildtype. Results from two independent *ifs1*Δ mutants are displayed, as well as from two spores that germinated from independent biological replicates of the *ifs1*Δ #2 mutant. All mutants were assayed in a set of at least 5 biological replicates alongside at least 5 wildtype replicates. Points display results from a single replicate, normalized to the mean from the corresponding wildtype controls. The boxplots summarize the underlying points and show first quartile, median, third quartile while the whiskers show the range of the data to a maximum of 1.5 times the interquartile range below and above the first and third quartile, respectively. Points outside the whiskers can be considered outliers.

**Figure 3 Supplement 4.**
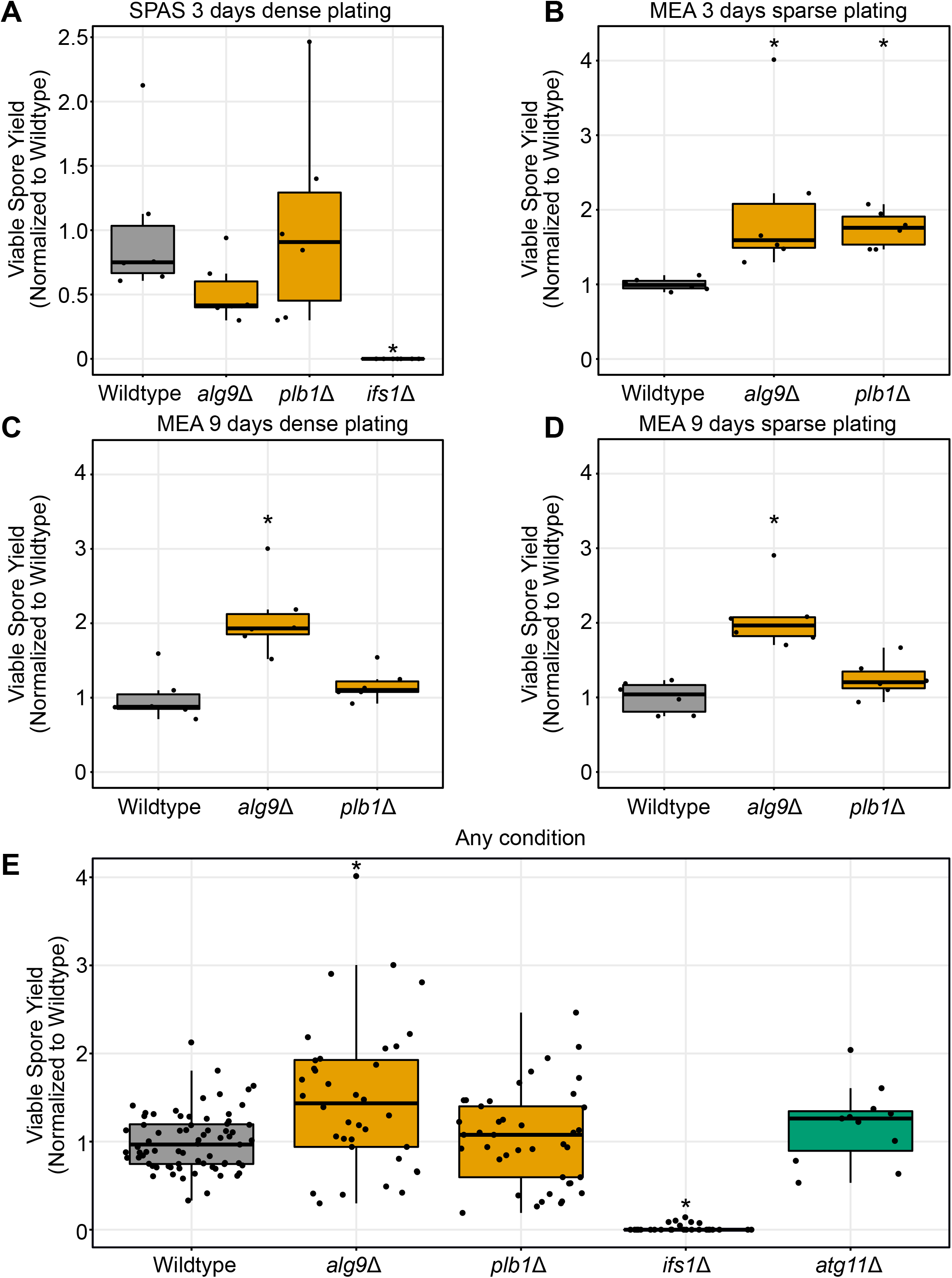
Viable spore yield varies by condition. A-E) Viable spore yield assay showing on the y-axis the number of spores produced per yeast cell plated, normalized to the mean value for wildtype. Points display normalized results from a single replicate. The boxplot summarizes the underlying points and show first quartile, median, third quartile while the whiskers show the range of the data to a maximum of 1.5 times the interquartile range below and above the first and third quartile, respectively. Points outside the whiskers can be considered outliers. A) Cells incubated on SPAS plates for three days at 25°C in dense cell patches (Mann-Whitney U test, *alg9*Δ, p=0.065; *plb1*Δ, p=0.94; *ifs1*Δ, p=0.0044). B) Cells incubated on MEA plates for three days at 25°C at low density as in the original TN-seq assay (Mann-Whitney U test, *alg9*Δ, p=0.0022; *plb1*Δ, p=0.0022). C) Cells incubated on MEA plates for nine days at 25°C in dense cell patches. (Mann-Whitney U test, *alg9*Δ, p=0.0043; *plb1*Δ, p=0.18) D) Cells incubated on MEA plates for nine days at 25°C at low density. These conditions match the original TN-seq assay, except that spores were not germinated in liquid. (Mann-Whitney U test, *alg9*Δ, p=0.0022; *plb1*Δ, p=0.24), E) Summary data encompassing all conditions tested for these three mutants. This includes data from Figure 3D as well as Figure 3 Supplement 4 A-D (Mann-Whitney U test, *alg9*Δ, p=0.0013; *plb1*Δ, p=0.73; *ifs1*Δ, p = 1.1*10^−14^; *atg11*Δ, p=0.14).

**Figure 4 Supplement 1.**
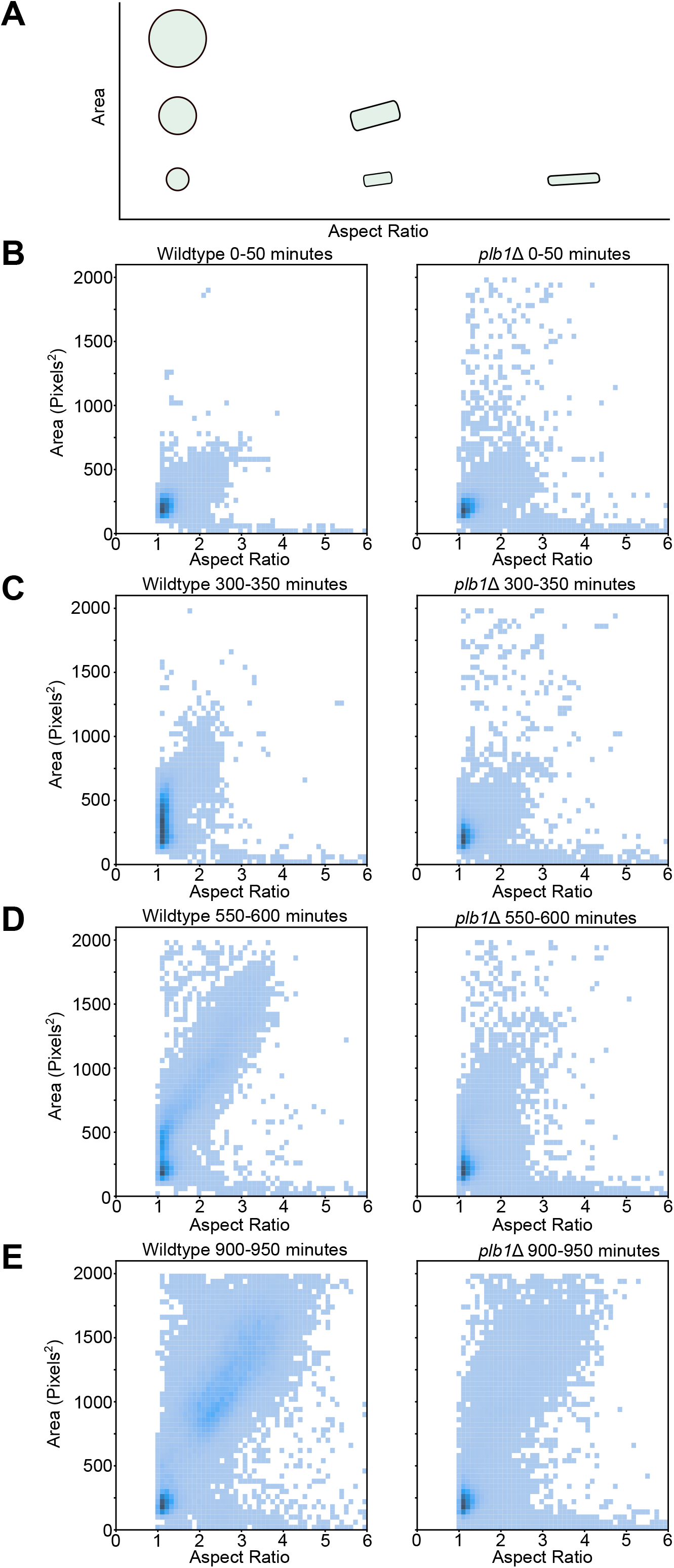
*plb1*Δ mutants exhibit a delay in growth and in establishing polarity. A) Schematic of plots of aspect ratio versus area. The x-axis displays aspect ratio, where 1 indicates round cells and larger numbers indicate more oblong cells. The y-axis displays spore size in square pixels. As shown in the cartoon, a higher aspect ratio indicates cells that are more elongated, while a higher area indicates cells that are larger. B-E) Two dimensional histograms showing the entire population of spores for wildtype or mutant in a given time window. B) Spores between 0 and 50 minutes after the video starts. C) Spores between 300 and 350 minutes. Wildtype spores have begun growing isotropically (i.e., swelling), but *plb1*Δ mutants have not. D) Spores between 550 and 600 minutes. Wildtype spores have continued to grow and begun to establish polarized growth. E) Spores between 900 and 950 minutes. Many wildtype spores have established polarized growth, while *plb1*Δ mutants are largely still small with only a subset establishing polarity.

**Figure 5 Supplement 1.**
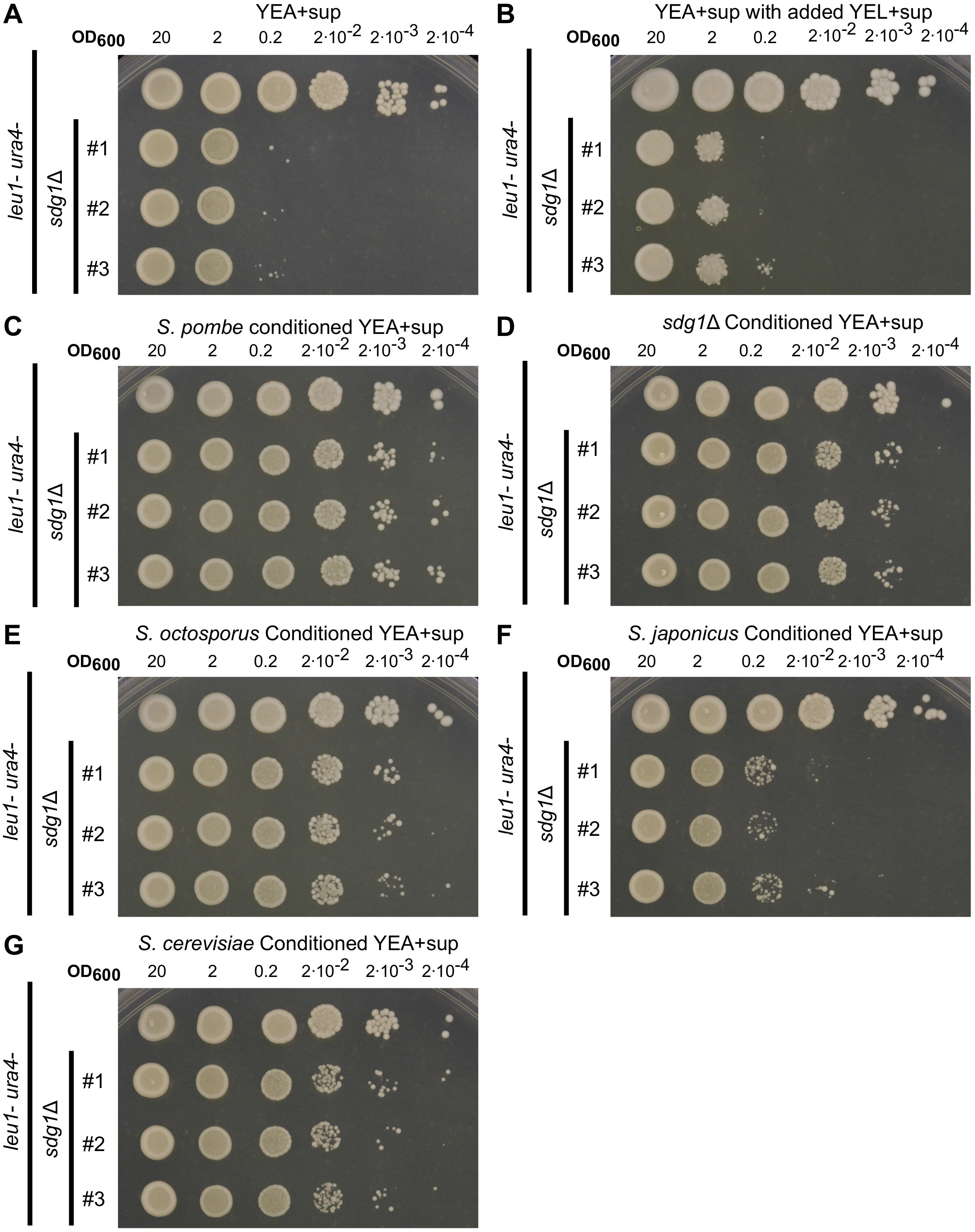
Conditioned media from different species provides varying levels of rescue for *sdg1*Δ mutants. A-G) Spot dilution assays with 5 μL spots plated. The initial leftmost spot is of OD600 = 20 culture and each successive spot is a 10-fold dilution, so that the final spot should be 10^5^ less concentrated than the first. All four experiments were conducted on the same day with the same dilution series of parent strain (*ura4*-D18, *leu1*-32) and three independent *sdg1*Δ mutants on the same genetic background (*sdg1*Δ::kanMX4, *ura4*-D18, *leu1*-32). All assays were also incubated for 4 days at 32°C. A) Spotted to standard yeast extract agar (YEA+sup) and. B) Spotted to YEA+sup where half the water had instead been replaced with yeast extract liquid with supplements (YEL+sup) media as a control for conditioned media. C) Spotted to conditioned YEA+sup media where half the water had instead been replaced with YEL+sup pregrown with the parent strain *S. pombe* (*ura4*-D18, *leu1*-32) (see methods). D) Spotted to conditioned YEA+sup media where half the water had been replaced with YEL+sup pregrown with an *sdg1*Δ mutant (*sdg1*Δ::kanMX4, *ura4*-D18, *leu1*-32). E) Spotted to conditioned YEA+sup media where half the water had been replaced with YEL+sup pregrown with *S. octosporus*. F) Spotted to conditioned YEA+sup media where half the water had been replaced with YEL+sup pregrown with *S. japonicus*. G) Spotted to conditioned YEA+sup media where half the water had been replaced with YEL+sup pregrown with *S. cerevisiae*.

**Figure 7 Supplement 1.**
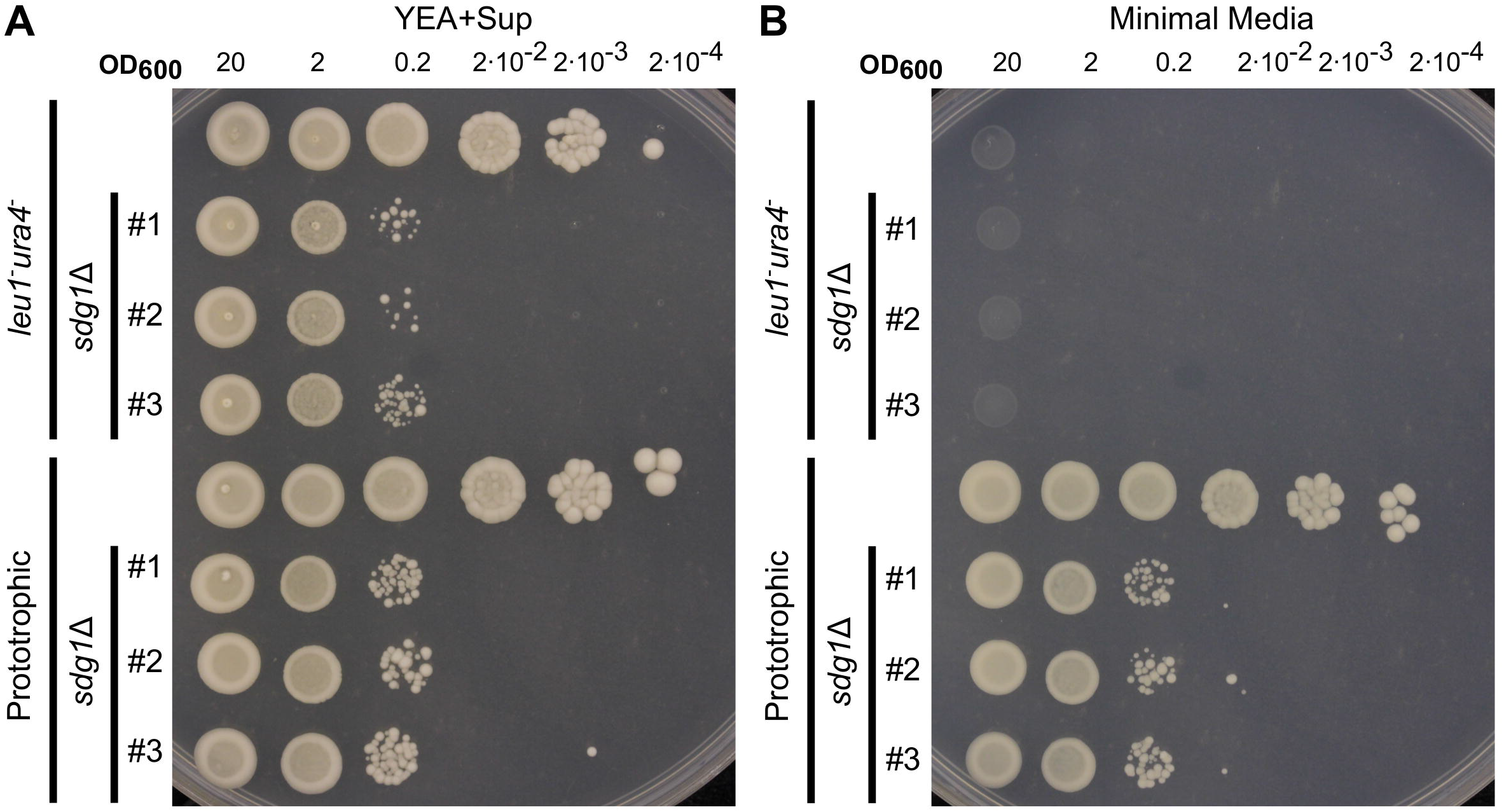
*sdg1*Δ does not cause auxotrophy. A-B) Spot dilution assays with 5 μL spots plated. The initial leftmost spot is of OD_600_ = 20 culture and each successive spot is a 10-fold dilution, so that the final spot should be 105 less concentrated than the first. Both experiments were conducted on the same day with the same dilution series of parent strain (*ura4*-D18, *leu1*-32), three independent *sdg1*Δ mutants on the same genetic background (*sdg1*Δ::kanMX4, *ura4*-D18, *leu1*-32), a wildtype prototroph (*h*^90^), and three independent prototrophic *sdg1*Δ mutants. A) Spotted to standard yeast extract agar (YEA+sup) and incubated for 4 days at 32°C. Note that the top half of this panel is the same experiment presented in Figure 5C. B) Spotted to minimal media and incubated for 4 days at 32°C.

**Figure 7 Supplement 2.**
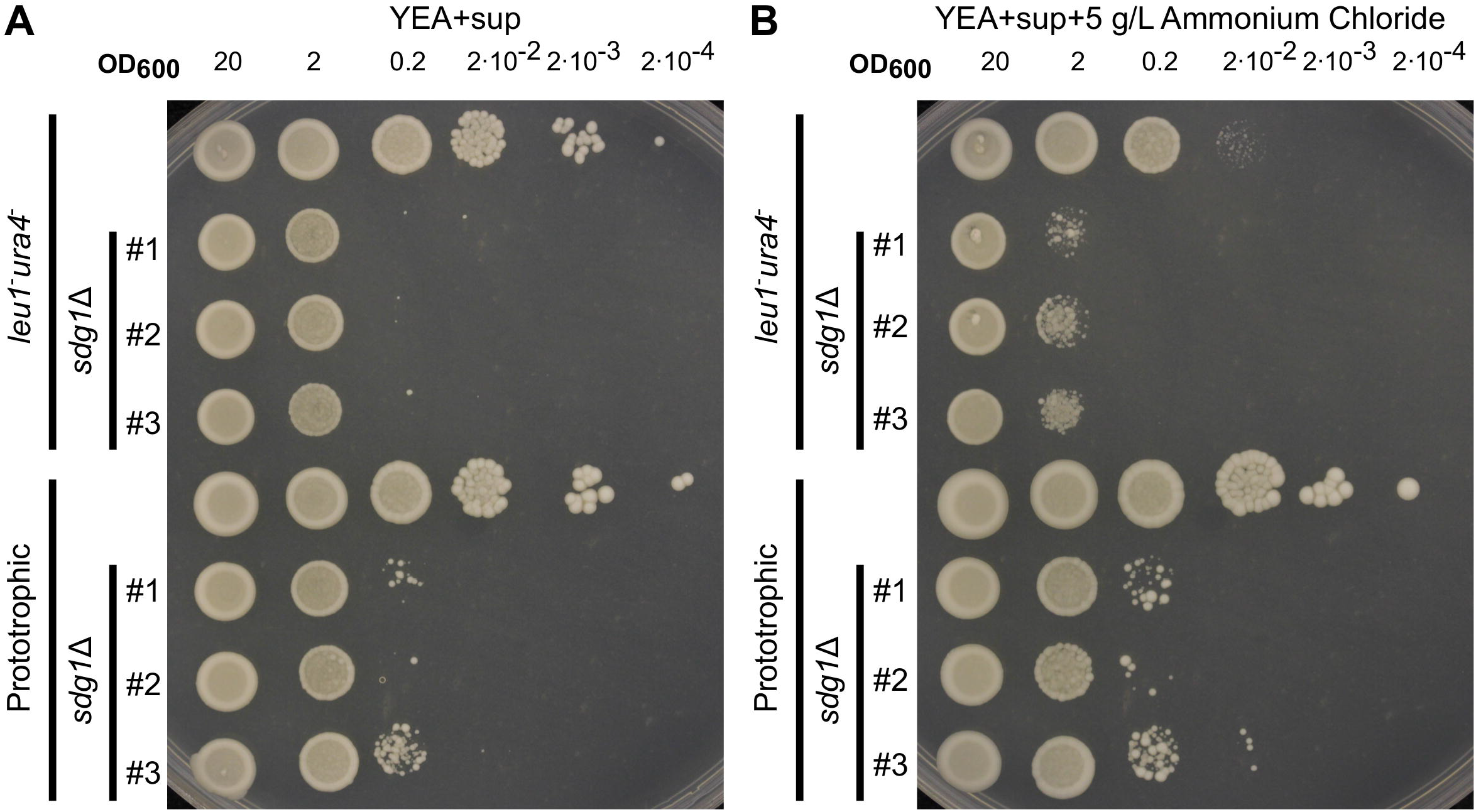
Ammonium chloride enhances low density growth defect of auxotrophic but not prototrophic *sdg1*Δ mutants. A-B) Spot dilution assays with 5 μL spots plated. The initial leftmost spot is of OD_600_ = 20 culture and each successive spot is a 10-fold dilution, so that the final spot should be 10^5^ less concentrated than the first. Both experiments were conducted on the same day with the same dilution series of parent strain (*ura4*-D18, *leu1*-32), three independent *sdg1*Δ mutants on the same genetic background (*sdg1*Δ::kanMX4, *ura4*-D18, *leu1*-32), a wildtype prototroph, and three independent prototrophic *sdg1*Δ mutants. A) Spotted to standard yeast extract agar (YEA+sup) and incubated for 4 days at 32°C. B) Spotted to yeast extract agar with 5 g/L supplemental ammonium chloride and incubated for 4 days at 32°C.

Table S1. Candidate Genes Identified by meiotic TN-seq

Table S2. Strains used in this study

Table S3. Oligonucleotides used in this study

Table S4. Plasmids used in this study

